# Nitroxoline treatment of *Balamuthia mandrillaris* triggers genomic crisis, transcriptional collapse, and incomplete encystment via copper and iron chelation

**DOI:** 10.1101/2025.02.25.640156

**Authors:** Kaitlin A. Marquis, Natasha Spottiswoode, Angela M. Detweiler, Norma Neff, Joseph L. DeRisi

## Abstract

*Balamuthia mandrillaris* is a free-living amoeba that can infect the central nervous system to cause granulomatous amebic encephalitis (GAE), a highly fatal disease with mortality rates up to 90%. GAE is typically treated with a six-drug regimen with limited efficacy and significant toxicity. Our lab previously identified nitroxoline, a quinolone antibiotic, as a promising therapeutic complement to the existing regimen, and in 2021 nitroxoline was first successfully used to treat a human patient. In this study, we elucidate nitroxoline’s mechanism of action using a combination of genomics, transcriptomics, chemical complementation, growth assays, and electron microscopy. In service of this goal, an annotated draft genome of *B. mandrillaris* strain CDC:V039 was generated using long read sequencing, yielding over 32,000 predicted protein coding sequences, the majority of which were present on telomere-to-telomere predicted chromosomal contigs. Leveraging this resource, comparative transcriptomics were used to characterize encystment responses to three different cellular stressors: nitroxoline, galactose, and hypoxia. Our findings reveal that nitroxoline-induced cysts undergo transcriptional collapse and genomic and structural destabilization via copper and iron chelation, and that preformed cysts are also disrupted by nitroxoline. These data provide insight into nitroxoline’s multifaceted impact on *B. mandrillaris* growth and cellular processes and advance our understanding of the transcriptional landscape associated with stress induced encystment. Our findings support the continued clinical study of nitroxoline as a complement to the existing regimen.

**Author Summary:** *Balamuthia mandrillaris,* one of three known brain-eating amoebas, causes a rare but usually fatal disease called granulomatous amoebic encephalitis. Until 2025, treatment options were limited, with most drugs proving ineffective against lab grown *Balamuthia* and toxic to humans. We previously discovered that the antibacterial drug nitroxoline prevented *Balamuthia* from destroying tissue by triggering encystment, whereby amoeba transition into dormant cysts. Nitroxoline was later used to successfully treat a patient, but it was unclear how the drug worked and whether the induced cysts could reawaken and cause the infection to return. To understand this process, we studied how *Balamuthia* responds to nitroxoline and compared it to two other stressors: low oxygen and excess sugar. Using genomic, chemical, and imaging tools, we found that *Balamuthia* activates different sets of genes depending on the type of stress, arguing against a single, universal encystment program. We also found that nitroxoline prevented formation of viable cysts and killed existing ones by depriving them of essential copper and metal ions. These findings improve our understanding of how *Balamuthia* becomes a cyst, offers clues for therapeutic intervention, and supports the continued use of nitroxoline to treat this deadly infection in combination with the existing drug regimen.

## Introduction

*Balamuthia mandrillaris* is one of three free-living amoeba (FLA) that are known to infect the central nervous system, along with *Naegleria fowleri* and *Acanthamoeba* spp. *B. mandrillaris* causes granulomatous amebic encephalitis (GAE) and skin lesions [1]. GAE due to *B. mandrillaris* is one of the single most morbid syndromes due to an infectious agent, with a mortality rate estimated at 90% in the United States [1,2]. Treatment for *B. mandrillaris* GAE has been hampered by limited understanding of the basic molecular mechanisms of the pathogen and by a lack of effective therapeutics.

Until 2024, the therapeutic regimen for *B. mandrillaris* GAE in the United States, according to the Centers for Disease Control and Prevention (CDC), [3] included six drugs: pentamidine, sulfadiazine, flucytosine, fluconazole, miltefosine, and azithromycin. This regimen is based on case reports of survivors [4,5], is inconsistently effective, and carries significant risk of toxicities. Of those six previously recommended agents, only pentamidine has an estimated 50% effective *in vitro* dose in the low micromolar range, [2,6–8] unfortunately; pentamidine is also highly toxic [9] and not thought to cross the blood-brain barrier effectively [10,11]. Mortality of patients who received at least one of the six standard medications remains very high, at an estimated 77% [2].

As an extremely rare and deadly infection, no randomized trials of possible therapeutic agents have ever been attempted. In the interests of improving the dismal survival rates from this infection, different groups have attempted to use *in vitro* techniques to identify other promising therapeutic agents. Studies have examined mold-active azoles (posaconazole, voriconazole or itraconazole) [7]; or used large-scale screens to identify other possible candidates such as the oncologic agent plicamycin (mithramycin) [12,13] and the tyrosine kinase inhibitor ponatinib [13]. In 2018, an unbiased growth inhibition screen of 2,177 drugs identified nitroxoline, a quinolone antibiotic historically used for treatment of urinary tract infections, as a promising *in vitro* candidate [6]. Nitroxoline is currently in use for urinary tract infections in some European countries and in China, and has a well-characterized and reassuring safety profile [14].

In 2021, a patient was diagnosed with *B. mandrillaris* GAE at the University of California San Francisco (UCSF). After the patient worsened clinically on the standard six-drug regimen, clinicians obtained and trialed the adjunctive use of nitroxoline, and the patient went on to survive his infection [15]. Since then, given the *in vitro* data and promising although limited clinical information, nitroxoline was added to the list of medications recommended to treat *B. mandrillaris* GAE by the Centers for Disease Control and Prevention (CDC). It is also now available to U.S. patients from the CDC through an expanded access Investigational New Drug protocol [16].

As nitroxoline gains increasing recognition as a possible addition to the regimen to treat *B. mandrillaris* GAE, a deeper understanding of the molecular effects of nitroxoline on this amoeba is warranted. Nitroxoline is a potent antibiotic with activity against a broad range of Gram negative and positive pathogens [17] and some fungi [18–20] by disrupting biofilm formation [21,22] and outer membrane integrity [23]. Mechanistic studies have revealed that nitroxoline chelates divalent metals including manganese (Mn^2+^) [24], magnesium (Mg^2+^) [24], iron (Fe^2+^) [22,25], copper (Cu^2+^) [19], and zinc (Zn^2+^) [22] in an organism specific manner. Previous structure-activity relationship experiments [6] with nitroxoline in *B. mandrillaris* suggested a similar mechanism of action, but the details of its impact on cellular functions are poorly understood, including which divalent cations may be involved.

*B. mandrillaris* has two life stages, including a motile trophozoite stage, and a tough, dormant cyst stage that is thought to be more resistant to drug treatment. Differentiation into cysts is stimulated in response to stress or harsh environmental conditions. Encystment can be experimentally stimulated by introducing a stressor to *B. mandrillaris* cultures, including noxious agents (galactose) [6,26] or other physiological stressors such as hypoxia. Cysts can then return to the motile trophozoite stage, a process termed recrudescence.

Nitroxoline is thought to induce amoeba encystment [6], but the details of encystment and amoeba response to this medication are poorly understood, including a basic understanding of transcriptomic changes relative to other stressors. Previous experiments indicated that nitroxoline treatment at low concentrations delayed recrudescence, while higher concentrations of nitroxoline prevented trophozoite recovery for a period of 28 days [6]. Together, this information suggested that nitroxoline may undermine the structural integrity of the cyst. While others have published transmission electron micrographs of *B. mandrillaris* cysts [27], neither the surface of the cyst nor the encystment process over time has been well characterized.

In this work, we address key questions regarding nitroxoline and its effects on *Balamuthia mandrillaris,* including a better understanding of the mechanism of action with respect to encystment, transcriptional integrity, and how nitroxoline impacts specific requirements of the amoeba for divalent metals. Through genomic, transcriptomic, and functional growth assays, supplemented by scanning electron microscope visualization, we shed light on these outstanding questions and inform future research and clinical directions.

## Results

To characterize the functional impact of nitroxoline treatment on *B. mandrillaris*, we undertook a comprehensive investigation that included transcriptomics, growth assays and chemical complementation, cyst integrity assessment, and inspection by scanning electron microscopy. To complete a transcriptomic study, accurate genome features, including gene models, predicted proteins together with predicted functions, complexes, and pathways are required for interpretability. Existing draft *B. mandrillaris* genome projects available through NCBI at the time of this writing were judged to be either too fragmented, and/or lacked publicly available gene model annotations [28–30]. Prior to this work, the most complete draft genome was derived from a different strain (ITSON01), [30] preliminary nucleic acid alignments suggested sufficiently diverse variation to warrant a genome assembly and annotation effort. Therefore, we undertook the production of a new draft genome for the lab strain of *B. mandrillaris* used in this study (V039) to specifically support our transcriptomic experiments. All genomic and transcriptomic data, as well as annotations, from this work will be publicly available at NCBI BioProject PRJNA1206197. Additional annotation data and an integrated browsing tool is available at: https://github.com/uc-derisilab/derisilab-standalone-balamuthia-browser.git

### Genomic DNA extraction and long read sequencing

High molecular weight genomic DNA (gDNA) was extracted from 100 million amoeba grown in 1,120 mL of axenic media, yielding 33.4 micrograms of gDNA with an average fragment size of 58 kb and DNA integrity score of 9, for PacBio HiFi long read sequencing. DNA was aliquoted into two separate halves; one half was sheared to produce ∼20 kb fragments in length, while the other half was size selected for an average fragment size of ∼60 kb (see methods for more detail). Two separate SMRT Cells 8M were sequenced on a PacBio Sequel IIe system (SMRTLink v. 11.0). The first SMRT Cell was used for the size selected gDNA fragments and yielded a total of 506,622 HiFi reads (Q36) comprising 5.9 Gb of sequence, with an average length of 11.6 kb. The second SMRT Cell was used for sheared gDNA fragments, and yielded 3,027,598 HiFi reads (Q38), comprising 22.2 Gb of sequence with an average length of 7.3 kb. Combining the data from both flow cells, this resulted in a pool of 3.5 million long reads and greater than 28 gigabases of sequence data.

Shotgun reads derived from eukaryotic genomes contain large numbers of mitochondrial reads. To create a consensus mitochondrial genome contig, reads greater than 41kb were screened by minimap2 [31,32] against previously sequenced *B. mandrillaris* mitochondrial DNA (mtDNA). A set of 19 such reads were assembled using the closest reference sequence (GenBank KT175741, strain V039) as a guide, which yielded a 40,046 nucleotide consensus sequence with a pairwise identity of 99.7% with the reference (NCBI BioProject PRJNA1206197). The consensus mtDNA sequence was further verified and confirmed using an independent collection of 2 million Q40 filtered paired-end (150x2) Illumina mtDNA reads mapped using STAR [33]. Using this new consensus mitochondrial genome sequence, the full collection of PacBio HiFi long reads were mapped back using minimap2. A total of 313,748 reads mapped to the mitochondrial genome, corresponding to 8.9% of the total reads and 6.7% of the total bases, suggesting a per cell mtDNA copy number of roughly ∼160, assuming an estimated genome size of 95 Mb.

### Genome assembly

High throughput chromatin conformation capture (Hi-C) has previously been used in conjunction with long read sequencing technology to produce high quality chromosome level assemblies for many organisms, including other free living amoeba species [34]. Five million amoeba were used to isolate 152 ng of purified DNA to construct Hi-C libraries. The final library yielded over 1x10^9^ raw reads (paired-end, 75 bp each). To create a high quality working subset, 100 million pairs of high quality full length reads (>Q30) were extracted using seqkit [35,36].

The overall assembly and annotation workflow is diagramed in S1 Fig. To create the input pool of reads for assembly, mitochondrial genome filtered PacBio HiFi reads were first sized selected to remove reads < 1kb, yielding 3.2 million reads total with an average length of 8.1kb. The de novo assembly tool hifiasm (v0.25.0-r726) [37,38] was used to create a draft. In a first pass, the input pool was error corrected using hifiasm default parameters, and then further size selected to retain reads > 10kb, yielding 663k reads with an average length of 13.9kb and a total size of 9.2 gigabases. A primary assembly was constructed with hifiasm using this size selected, error corrected input pool, leveraging the collection of 100 million quality filtered HiC read pairs, using default parameters. Previous inspection of raw sequence reads revealed that the *Balamuthia* telomeres are of the canonical repeat “TTAGGG” variety, and thus the “—telo m” hifiasm option with this sequence was specified using this sequence. The primary assembly consisted of 308 contigs. Inspection revealed 102 contigs contained telomeric repeats at both ends, suggesting these were complete, telomere-to-telomere (T2T) contigs.

Analysis of k-mers with the meryl/merqury package (1.4.1) [39] provides an objective measure of completeness and quality by comparing the assembly to the input read pool in both T2T contigs and loose contigs. Using a kmer size of 21, contigs contributing zero or less than 0.1% of all k-mers were removed, resulting in a total of 96 loose contigs in addition to the 102 T2T contigs. To scaffold the remaining non-T2T contigs, the YaHS [40] scaffolding tool was used with the 100 million HiC read pairs as input. Redundant contigs were eliminated from the output and combining the results yielded a total of 186 contigs, of which 113 were T2T contigs and 73 were non-T2T contigs. Of the 73 non-T2T contigs, 53 (72%) feature telomeric repeats on one end of the contig. Merqury analysis using the input pool of (>10kb) HiFi reads resulted in a completeness score of 99.3, and an exceptional QV score of 56.1 indicating excellent consensus base accuracy. The total length of this draft assembly is 94.77Mb. Contigs were renumbered 001 to 186, sorted largest to smallest, with a prefix “T2T” indicating telomeric repeats at both ends, or “SCF” for contigs with one, or no telomeric repeats. The T2T contigs comprise 66.3Mb (69.9% of the total). The distribution of contig sizes ranged from the largest, T2T001, with a length of 1.49Mb, to the smallest (SCF186) at 43.3kb (S2 Fig). The average length of telomeric repeats was 540bp or 90 repeats. Genome data for this project is available at NCBI BioProject PRJNA1206197.

### Gene models and annotation

Prior to predicting gene models, tRNA-Scan-SE (v2.0) [41], Infernal (v1.1.5) [42], EDTA (v2.2.2) [43], and Dustmasker (v1.0.0) [44] were used to annotate non protein coding regions of the draft genome assembly including tRNA, ribosomal and non-coding RNAs, repeat regions and transposable elements, and low complexity regions, respectively. A total of 681 tRNAs, 389 ribosomal RNAs and 100 non-coding RNAs were annotated with tRNA-Scan-SE and Infernal (Fig 1). Approximately 13.37% of the genome was found to be comprised of low complexity regions by Dustmasker and up to 16.04% of the genome were annotated as transposable elements/repeat regions by EDTA at the maximal sensitivity threshold.

**Fig 1.**
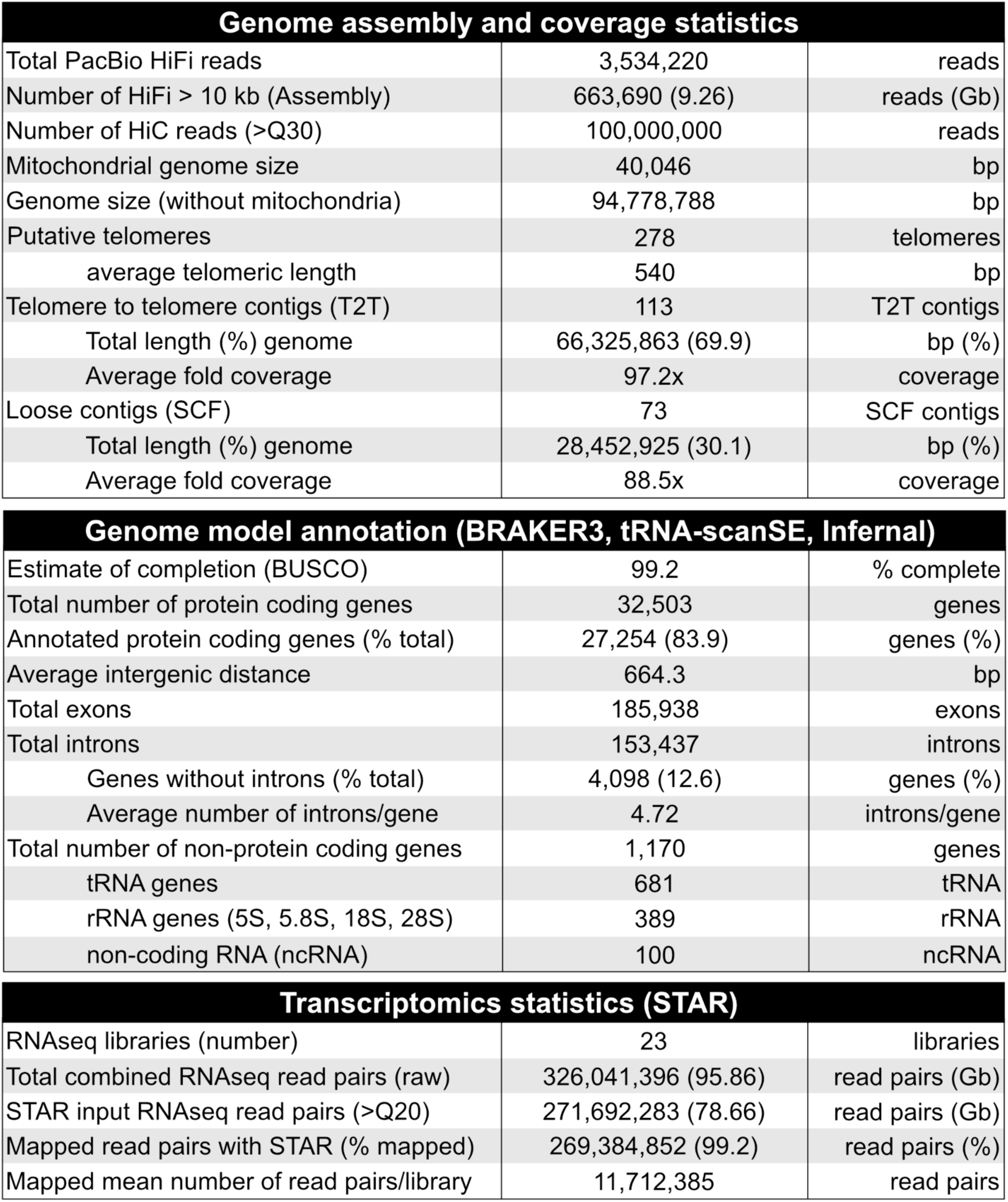
*Balamuthia mandrillaris* UCSF CDC:V039 genome model summary. Summary of genome assembly, annotation, and transcriptomic statistics for *Balamuthia mandrillaris* UCSF CDC:V039 strain. Metrics include sequencing depth, assembly completeness, gene model annotation, and RNA-seq mapping statistics.

After annotating these regions, four different gene models were created using the BRAKER3 package [45]

with varying sensitivity thresholds for soft masking (S1 Appendix) using our draft genome assembly and the complete set of RNA sequencing data (23 independent libraries, 326 million read pairs total), described in detail below, as input. Given that gene model predictions often varied, every prediction was manually curated using Integrative Genomics Viewer (IGV, version 2.19.6) [46]. If all four BRAKER3 predictions were inconsistent with the RNAseq and splice junction data, gene models were manually corrected using Geneious Prime (v2025.0.3). After curation, a total of 32,503 protein coding genes were annotated (Fig 1). This draft annotation consists of 32,503 putative protein coding sequences, featuring an average of 4.7 introns per transcript. Introns are relatively short, averaging 113nt, and less than 13% of predicted coding sequences were without any introns. The most highly spliced transcript in the genome (belonging to T2T133g112675) has 63 introns with a distinctive repeating exon structure which possess similarity to ice nucleation repeats found in prokaryotes. The average intergenic distance in this draft is approximately 664bp, consistent with a highly compact genome with a protein coding content of greater than 50%. Although a strict comparison is not within the scope of this work, we note that the set of predicted proteins from the strain of *Balamuthia* used in this study (V039) and those from the previously sequenced ITSON01 strain had a mean normalized identity of approximately 90%, in an all-by-all blastp analysis (v2.12.0). Approximately 5% of proteins did not yield a match, suggesting differences in assembly (fragmentation), gene models, completeness, or true divergence.

We also note that this genome is replete with putative transposable elements (TEs) or portions thereof. While most of these are captured by this annotation pipeline, the highly fragmented nature of many of these elements poses an annotation challenge. A thorough annotation of TEs will require a future dedicated effort to be comprehensive.

Assessment using BUSCO (v6.0.0) [47,48] in protein mode revealed a completeness score of 99.2%, suggesting that a near full complement of the expected eukaryotic proteins is present in this assembly. The single missing BUSCO was phosphoacetylglucosamine mutase (PGM3, OrthoDB 1928at2759). While highly confident predictions for the enzymatically similar phosphoglucomutase PGM2 were evident (T2T138g113305 and SCF153g157325), no credible predicted protein for PGM3 was detectable using additional search methods. We note that an all-versus-all blastp analysis with reciprocal best-hit filtering identified one or more high-identity counterparts for more than 94% of predicted sequences, while no credible counterparts could be determined for the remaining 6%. Only 33% of proteins yielded a single counterpart, suggesting widespread gene family expansions and may explain non-uniform correspondence between putative homologous contigs. The BUSCO analysis indicated that core conserved proteins are almost all duplicated (95%) supporting strong diploid-like genome character, but with apparent structural heterogeneity, uneven haplotype correspondence, rearrangements, gene-content heterogeneity, and non-uniform collinearity of proteins among many putative homologous contigs. While this complex genome will require further structural characterization, completeness metrics suggest the complete, or nearly complete repertoire of coding sequences is captured here.

To produce GO terms and predicted gene functions, a multi-pronged functional annotation strategy was used, combining the aggregated results produced by the InterProScan Suite of analysis tools (5.77) [49], ProteInfer [50], PANNZER-2 [51], eggNOG-mapper v2 [52], and protein similarity using DIAMOND [53] to query the NCBI NR database [54]. The complete set of gene models, corresponding protein sequences, and annotations are available as part of NIH BioProject PRJNA1206197 and through our searchable, standalone genome browser.

### The transcriptional landscape of stress induced encystment in *Balamuthia mandrillaris*

Previous work has shown that nitroxoline provokes encystment of *B. mandrillaris*, presumably as a stress response [6]. Therefore, a characterization of encystment in the absence of drug is also necessary, similar to what has been accomplished for *Acanthamoeba* [55], another free-living amoeba that infects the human brain. To this end, *B. mandrillaris* trophozoites were stimulated to encyst by three distinct perturbations: with 12.5 μM nitroxoline, 12% galactose, or with an unrelated cellular stress, namely microaerophilic gas generating sachets (hypoxia). These experiments, to our knowledge, are the first such experiments explicitly measuring time courses of transcriptional change in *Balamuthia* during axenic growth, galactose, nitroxoline, and hypoxia. RNA was isolated prior to drug treatment, and then at 8, 24, 48, 72, and 168 hours (7 days) post treatment (Fig 2A). Phase contrast images were taken at the time of RNA isolation to verify encystment status (S3 Fig). Nitroxoline, galactose, and hypoxia samples all showed obvious signs of cell rounding by 24 hours, with most of the population consisting of rounded cyst-like morphologies by 48 hours. Nitroxoline and galactose induced cysts did not recrudesce during the entire 7-day treatment period. In contrast, hypoxia treated cells appeared to adapt to the low oxygen environment and rapidly transitioned in and out of cyst-like morphologies by 72 hours post treatment. At the end of the 7-day treatment course, galactose cultures were comprised of essentially 100% cysts while nitroxoline treated samples appeared cyst-like but degraded (S3 Fig). Furthermore, no intact RNA could be isolated from the 7-day nitroxoline treated amoeba, therefore this time point was excluded from further analysis.

**Fig 2.**
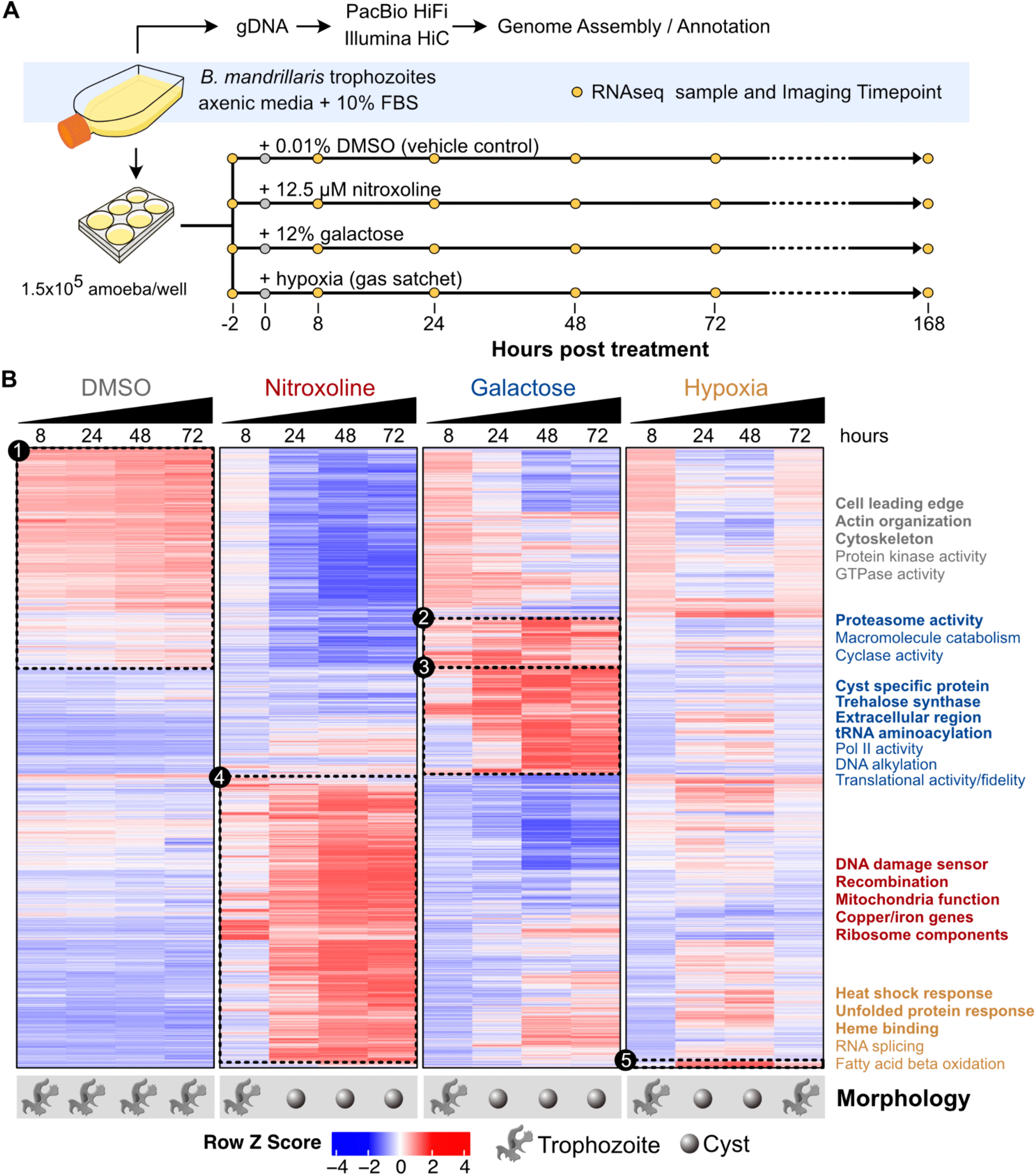
Transcriptional responses to stress induced encystment in *Balamuthia mandrillaris.* (A) Overview of experimental design. Genomic DNA (gDNA) was isolated from axenically grown trophozoites, sequenced, and assembled into an annotated draft genome to support transcriptomics. Stress induced encystment was triggered in trophozoites with nitroxoline, galactose, or hypoxia and RNA was extracted at indicated times post treatment. Amoeba morphology was examined microscopically at the time of extraction (S3 Fig). (B) Heatmap of 12,328 differentially expressed genes from stress induced treatments over time (168-hour timepoint excluded due to degraded RNA). Five expression clusters of interest are highlighted with enriched genes and/or gene ontology terms (S4 Fig, S1 Table).

Transcriptional changes in response to different cellular stressors were characterized using the DESeq2 package from R [56] by collapsing time point samples into biological replicates and computing regularized log transformed counts for differential expression. Using a threshold value of log2 fold change greater than 1 and adjusted p-value less than 0.01, we identified 12,328 genes with significantly different expression between encystment conditions. This set of genes were hierarchically clustered using complete linkage Pearson correlation [57], converted to row z scores, and visualized in a heatmap (Fig 2B, S1 Table). Five dominant gene expression clusters that visually discriminated between sample treatment conditions were extracted for gene ontology (GO term) enrichment using the topGO R package [58,59] and aggregated genome annotations (Fig 2B, S1 Table, S4 Fig). A standalone genome browser, featuring searchable annotations, expression data, and gene models with coverage plots was constructed to aid interpretation and exploration of the data, and is available at https://github.com/uc-derisilab/derisilab-standalone-balamuthia-browser.git.

Expression cluster 1, which contains 3,378 predicted genes, was characterized by upregulation in DMSO treated trophozoites and was dominated by genes primarily involved in general cell growth and locomotion. Enriched GO terms included genes associated with the leading edge of the cell, including actin and cytoskeletal organization. Actin itself is highly represented in the genome, numbering 26 identical copies, all in different genomic loci, and all predicted as single exon genes, devoid of introns. In the unperturbed DMSO control condition, instances of actin were also among the genes with the highest expression, often exceeding thousands of transcripts per million (TPM). Examples of actin related functions include SCF042g124140 and T2T026g037995, which encode putative alpha and beta components of F-actin capping protein complex respectively, a key function for filament dynamics, stable cell migration and motility. Likewise, numerous putative components of the Arp2/3 complex (S1 Table), critical for cellular growth, filamentous actin, and polarized migration, are also represented, presumably involved in pseudopodia formation observed in trophozoites. Additional hallmarks of actin and cytoskeletal engagement include tropomodulin T2T003g007645, T2T002g004960, villin-like protein (actin severing) T2T004g009135, and several putative myosins such as T2T019g028490 and T2T018g027270. Many of the individual genes from cluster 1 were also predicted to contain protein kinase and GTPase activity, which are important effector intracellular signaling domains regulating cell cycle, cell growth, differentiation, morphology, and movement [60]. Other notable genes in this cluster included proteins with predicted roles in DNA replication, transcription, translation, and mitotic regulation. Notably, the corresponding genes in the nitroxoline treated cells are uniformly down regulated, consistent with the cessation of growth and motility cessation observed after drug treatment.

Cluster 2 is an overlapping subset of Cluster 1 and contains over 990 predicted coding sequences that were upregulated similarly in both DMSO treated trophozoites and galactose induced cysts, with slightly higher expression overall in galactose, while the corresponding genes in nitroxoline treated amoeba were strongly down regulated. Cluster 2 was enriched for general GO terms involved in intracellular signaling and cell homeostasis (modification-dependent protein catabolism, regulation of metabolic process, phosphorelay sensor kinase activity). Molecularly, these genes were enriched for threonine-type endopeptidase activity, proteasome activating activity, and cyclase activity. This cluster was also highly enriched for predicted proteins relating to protein turnover. This includes numerous putative subunits of the 26S proteosome itself (S4 Fig), including the 19S regulatory complex, for example SCF032g122040 (20S alpha-3), T2T081g085000 (20S alpha-5), and T2T038g046545 (20S beta-2), and 19S subunits such as SCF028g119880 (RPN8-like), T2T099g095665 (RPN2-like), and T2T014g022730 (RPT6a-like) and other associated degradation machinery (ie. E3 ubiquitin protein ligases, such as the HECT domain UPL1-like protein T2T003g007575, and the UBA52-like L40-Ubiquitin fusion protein T2T002g003505), and autophagy components (S1 Table). Together, the putative processes represented in cluster 2, which represents the overlap between Cluster 1 (rapid growth), and galactose induced encystment are consistent with processes associated with normal cellular growth, protein turnover/degradation, and core metabolic functions (S1 Table).

Cluster 3 comprises a large set (2,128) of predicted genes that were differentially enriched in amoebas induced to encyst by galactose treatment. These induced cysts were uniquely enriched for GO terms associated with the extracellular region, tRNA aminoacylation and ligase activity, enzymes related to carbohydrate storage including trehalose and glycogen, and multiple close relatives of cyst specific proteins (UspA, universal stress protein-like) in *Acanthamoeba* (ie. T2T077g082940, T2T092g090915, T2T012g020125) (S4 Fig, S1 Table). Genes belonging to Cluster 3 also include nearly a full complement of putative specific tRNA ligases, tRNA synthetases, tRNA hydrolases, and associated processing components, including multiple instances of the tRNA processing endonuclease RNAse Z (for example T2T039g047840, T2T075g080615, T2T102g097585, SCF139g151445). Other cluster 3 genes include those related to dauer up-regulated-related proteins (SCF136g149935, T2T045g053580), which are known to confer resistance to desiccation in *C. elegans* [61]. Gene members of Cluster 3 may also belong to specialized versions of core growth proteins. One example is translation elongation factor 1-alpha (eIF1alpha) for which there are at least 8 paralogs annotated in this assembly, forming two distinct groups (S1 Appendix). One group of eIF1alpha paralogs was uniformly expressed in all conditions except galactose induced encystment, where they were diminished. The other group were dramatically upregulated during galactose induced encystment, often greater than 1000-fold by TPM. Within each group, the proteins are essentially identical, but between the two groups, there is only ∼64% pairwise protein identity, suggesting the possibility that *Balamuthia* utilizes specialized paralogs during encystment for selective mRNA translation or altered translation dynamics.

Cluster 4 is the largest of any of the expression clusters with over 5,500 differentially expressed genes (Fig 2B) and contains genes that are significantly upregulated after treatment with nitroxoline. While nitroxoline treatment appears to morphologically initiate encystment, the transcriptional profile is distinct from galactose induced cysts. An overwhelming majority of protein annotations and GO terms in this cluster were directly related to mitochondria structure or function (mtDNA encoded proteins, protein targeting to mitochondrion, mitochondrial protein-containing complex, mitochondrial ribosome, and mitochondrial inner membrane), and DNA repair. Other enriched GO terms included respiratory chain complex IV assembly, many structural constituents of ribosome (for example, L15, L17, L30, L24, L29, L34, S10, S12, S14, S19, S26, and many more), and ATP-dependent DNA damage sensors, sterol and lipid-related enzymes (S4 Fig). Individual genes within cluster 4 include numerous putative components of DNA damage repair pathways, including canonical repair proteins, DNA damage checkpoint proteins, and DNA damage sensors. The DNA damage response is described in additional depth in the section “Nitroxoline results in rapid induction of the DNA damage response” that follows. DNA repair adjacent proteins were also enriched for in Cluster 4, including histone H2B, histone H2-like proteins, and histone deacetylase complex subunits. Of note, cluster 1 genes, associated with normal proliferative growth, are significantly down regulated in response in nitroxoline (Fig 2, S1 Table). Taken together, transcriptional response to nitroxoline treatment bears little similarity to galactose induced encystment and is suggestive of a cessation of normal proliferative growth, mitochondrial crisis, and a DNA damage dominated stress response.

Lastly, gene cluster 5 is a small subset of 135 genes that were highly upregulated in hypoxia induced cysts across all treatment timepoints and may represent genes involved in adaptation to the low oxygen environment. Two of the most overrepresented GO terms in this cluster were cellular heat acclimation and unfolded protein binding, suggestive of a hypoxia-induced heat shock response (S4 Fig). Specifically, Hsp20 (T2T021g031225), Hsp70 (T2T114g106765), and Hsp90 (SCF006g117605) family members (S1 Table) were all highly expressed in this cluster. These proteins are thought to play critical roles in the ability of cells to adapt to low oxygen environments [62,63]. Other overrepresented GO terms in this cluster included genes involved in splicing (spliceosomal complex, U1 snRNP, and mRNA splice site recognition).

### *Balamuthia mandrillaris* response to nitroxoline induces expression of genes encoding metal binding proteins

Given nitroxoline’s well documented metal chelating ability, the expression of genes with putative functions related to metal binding and/or utilization were interrogated, with a particular focus on copper, iron, and zinc - the three most abundant trace metals in the human brain [64]. Metal-related genes were queried using a list of copper, iron, and zinc associated GO terms (S1 Appendix) manually curated from QuickGO [65]. Likely paralogs were collapsed and averaged based on a DIAMOND normalized bit score of 0.98. Regularized log transformed counts from DESeq2 were averaged for identical genes yielding a set of 1,405 genes encoding putative proteins associated with copper, iron, and/or zinc. Genes were binned by transcriptional similarity, yielding a single cluster of 123 unique genes that were highly upregulated in response to nitroxoline (Fig 3A, S2 Table). The overall gene expression pattern for the cluster was summarized by summing the row Z scores of each category contained within (Fig 3A -“Z score Sum”). Consistent with nitroxoline acting as a divalent chelator of metals, genes associated with copper and iron metabolism were upregulated in response to nitroxoline, including iron sulfur cluster assembly genes (NiFU-like, NBP35-like, NUBP1, NUBP2) and copper homeostasis and chaperone proteins (CutC-like, COX11, COX19, SCO1) (S2 Table). This cluster was also enriched for several mitochondrial proteins, notably mitochondrial iron sulfur cluster assembly and trafficking proteins (NFS1 and GLRX5) and proteins involved in mitochondrial transport and metabolism (LIAS and MDL1). Other notable upregulated genes involved those associated with controlling the cellular redox state (FDX2 and superoxide dismutase) as well as DNA damage and repair associated enzymes (PARP, ALKBH3, RFC4, and DNA glycosylase), as discussed below.

**Fig 3.**
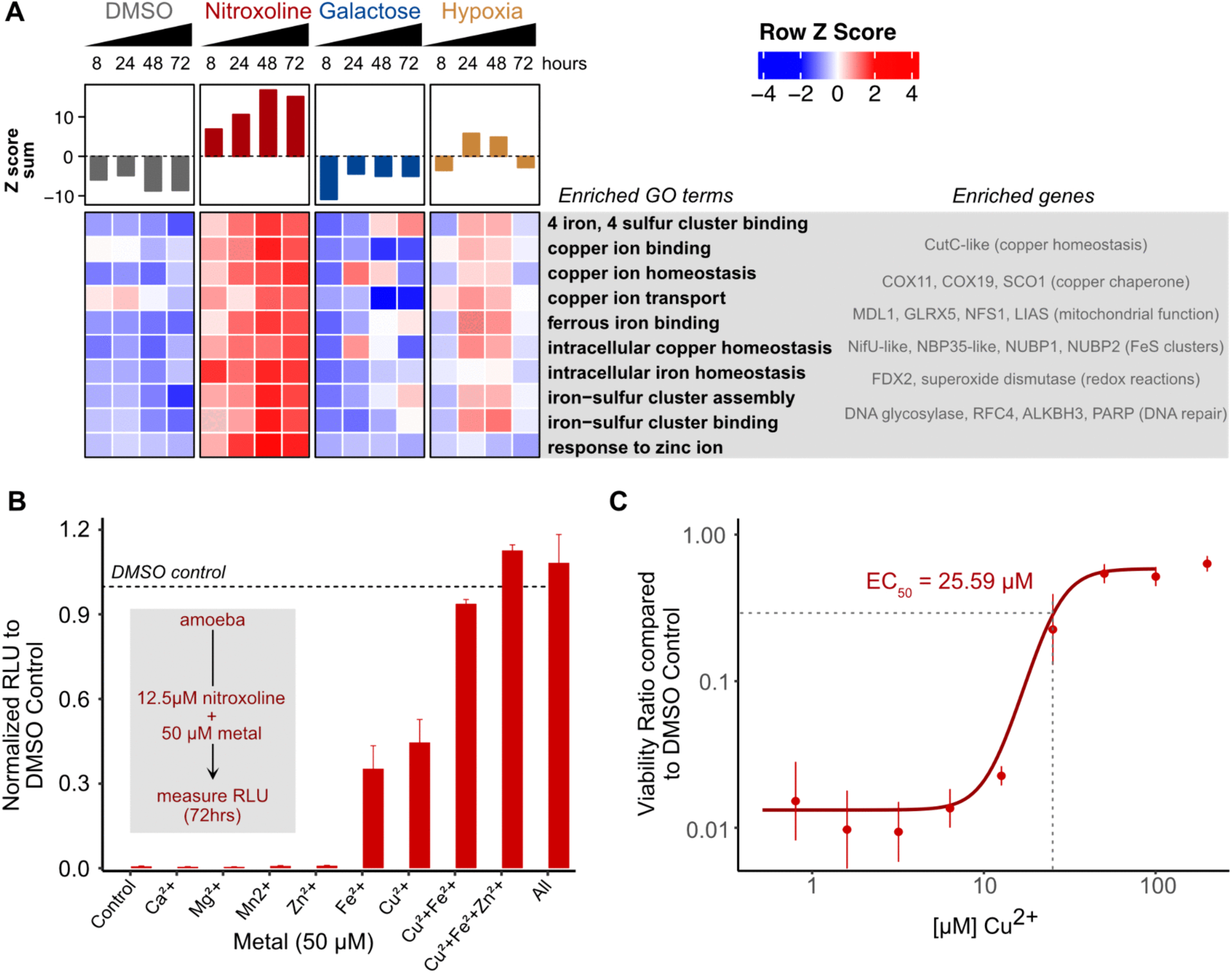
Nitroxoline inhibits *B. mandrillaris* by disrupting metal-associated gene expression via copper and iron chelation, with rescue by exogeneous metal supplementation. (A) Cluster of 123 unique copper, iron, and zinc associated genes that were upregulated in response to nitroxoline treatment. Metal associated genes were identified using a list of over 50 manually curated GO terms (S1 Appendix) and hierarchically clustered by expression. Proteins sharing more than a normalized bit score of 0.98 (by DIAMOND) were averaged prior to computing individual row Z scores. All row Z scores were summed to provide an overall expression profile. (B) Exogenous supplementation with copper, iron, and zinc maximally rescues nitroxoline mediated inhibition. Divalent calcium (Ca2+), magnesium (Mg2+), manganese (Mn2+), zinc (Zn2+), iron (Fe2+), and/or copper (Cu2+) were incubated with nitroxoline or DMSO and viability was assessed 72 hours later using a luciferase-based assay. Relative light units (RLU) were normalized to respective DMSO controls (n = 3). (C) Copper rescues nitroxoline mediated inhibition at a stochiometry ratio of 2:1. Amoeba were treated with 12.5 µM nitroxoline or DMSO control with increasing concentrations of copper for 72 hours. Viability was measured with luciferase assay and normalized to respective DMSO controls (n = 3).

### Nitroxoline induced killing of *Balamuthia mandrillaris* is rescued by supplementation with copper and iron

Nitroxoline’s ability to chelate divalent metals is well known [24,66,67], however it is not known if metal chelation is related to the nitroxoline induced killing of *Balamuthia*. To investigate the role of divalent metals relative to growth inhibition by nitroxoline treatment, viability assays were conducted with exogenous divalent metal supplementation. Amoeba were treated with 12.5 μM nitroxoline and supplemented with 50 μM of divalent cations. These concentrations were chosen based on morphological screening of cultures with varying amounts of nitroxoline and metal to minimize cytotoxic effects. Under these treatment conditions, iron and copper each partially rescued viability and prevented morphological encystment, with copper demonstrating a slightly superior rescue (Fig 3B). Co-supplementation with copper and iron resulted in essentially 100% rescue. Co-supplementation with copper, iron, and zinc resulted in a full rescue, slightly greater than that of vehicle controls, and recapitulated the effects of all trace metals supplemented at once. The dose response to increasing amounts of supplemental copper in the presence of 12.5 µM nitroxoline revealed an EC50 of 25.5 µM (Fig 3C). The IC50 for nitroxoline is 3-4 µM for *in vitro* growth inhibition [6]. Interestingly, the FDA-approved iron chelator deferoxamine resulted in a relatively poor 157.3 µM IC50 (S5 Fig) suggesting drug influx/efflux or intracellular action differs between these compounds. These data are consistent with the known chelation role of nitroxoline, with respect to iron and copper specifically.

### Nitroxoline results in rapid induction of the DNA damage response

Previous reports suggested that nitroxoline induces oxidative stress and chromatin condensation in *Naegleria fowleri* [68] and the expression of several DNA repair transcripts in *Acanthamoeba castellani* [69]. Here, treatment of *B. mandrillaris* with nitroxoline also yielded a rapid and marked signature of expression induction in DNA repair. Employing the same annotation strategy as described above, 329 unique genes were identified belonging to 16 different DNA damage and repair categories (Fig 4A, S3 Table, S1 Appendix). This gene set spans the entire gamut of canonical DNA repair pathways, including base excision repair proteins (NEIL1, UDG, AP endonuclease); nucleotide excision repair factors (ERCC1, XPC, XPG); mismatch repair components (MutS and MutL homologs); the double-strand break repair protein MRE11; non-homologous end joining proteins (NHEJ1, DNA Ligase IV, XRCC4); homologous recombination components (RAD51, BRCA2-like); checkpoint repair proteins (HUS1, RAD9B, RAD1); and Fanconi anemia pathway proteins (FANCA, FANCI, FANCL, FANCD2, FANCM) involved in interstrand cross-link repair, consistent with widespread activation of genomic damage pathways. Beyond these core pathway components, a broader set of putative DNA repair orthologues were induced by nitroxoline treatment, including putative exonucleases, helicases (Mcm8/9), PCNA, dealkylating enzymes, telomere regulation factors, Spo11-like proteins, TatD-like DNA hydrolyases, and many others (Fig 4A, S3 Table). Increases in expression of DNA damage and repair associated genes were evident as early as 8 hours post treatment with nitroxoline, preceding the formation of obvious cysts (S3 Fig), and differentiating this response from galactose induced encystment. These pathways remain elevated over the duration of the experimental time course in nitroxoline treated samples, whereas only a modest increase of these pathways is observed at 48 and 72 hours post treatment with galactose. No increase in DNA damage associated gene expression was observed in hypoxia or DMSO treated samples.

**Fig 4.**
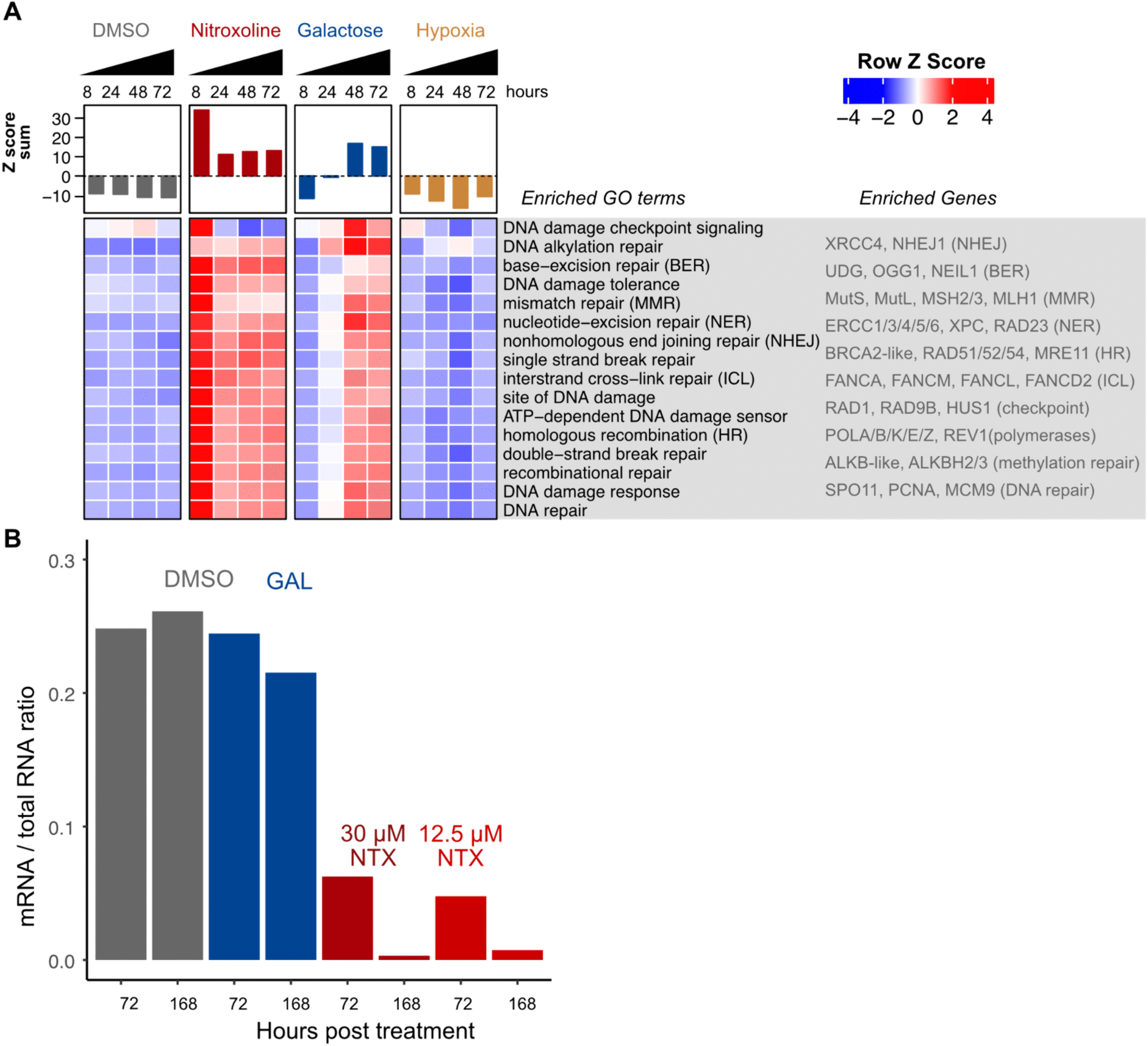
Nitroxoline induces genomic stress and perturbs global mRNA transcription. (A) Nitroxoline induces early and excessive expression of DNA damage and repair associated genes at 8 hours post treatment. Expression patterns for 329 unique genes were collapsed into enriched gene categories using manually curated GO terms (S1 Appendix). Each category contains at least 10 unique genes, and all row z scores were summed to provide overall expression profiles. (B) Global mRNA transcription is impaired in nitroxoline induced encystment. Encystment was stimulated with 12% galactose (GAL), 12.5 μM, or 30 μM of nitroxoline (NTX) for 72 and 168 hours and compared to DMSO treated trophozoites. Total RNA was extracted, and the proportion of short-lived mRNA reads were normalized to total RNA reads for each sample.

### Nitroxoline induced DNA damage is associated with loss of transcriptional integrity

Genome integrity, the response to DNA damage, and mRNA transcription are intimately linked cellular processes [70]. Widespread genome damage would be predicted to disrupt normal transcriptional activity; therefore, we quantified the proportion of short-lived messenger RNA (mRNA) relative to total RNA (dominated by long lived rRNA species). Nitroxoline was tested at two different concentrations, 12.5 μM and 30 μM, which prevents recrudescence of trophozoites in a 28-day time course [6]. Total RNA was extracted at 72 hours and 168 hours post treatment with nitroxoline, galactose, or DMSO treated samples and RNA fractions were sequenced using random primers (to capture mRNA and non-mRNA) and mapped back to our annotated genome using STAR [33]. DMSO treated trophozoites and galactose induced cysts (Fig 4B) yielded a nearly identical ratio of 0.25 (mRNA/total RNA). In contrast, nitroxoline essentially abolished the fraction of mRNA at both 72 and 168 hours, consistent with catastrophic cessation of normal mRNA production and/or increased mRNA degradation.

### Nitroxoline does not induce encystment associated marker genes

Nitroxoline appears to initiate encystment morphologically (S3 Fig), but the overall transcriptional program is significantly different than either galactose or hypoxia (Fig 2B). In *Acanthamoeba* and other amoeba, several cyst-specific proteins have been previously identified, including the aptly named “cyst specific protein” and proteins associated with trehalose production [55,71,72]. To further investigate encystment in response to nitroxoline, galactose, and hypoxia, the transcriptional response of *B. mandrillaris* genes encoding putative cyst specific proteins and trehalose pathway members were compared.

Trehalose synthetase, the first and rate-limiting enzyme in trehalose biosynthesis, catalyzes a critical step in producing this protective disaccharide, which stabilizes proteins and confers long term resistance to desiccation and other cellular stressors in a wide range of organisms, including amoeba [72–77]. The two key proteins associated with trehalose biosynthesis, trehalose-6-phosphate synthase (TPS) and trehalose-6-phosphate-phosphatase (TPP), were identified using InterPro accessions IPR003337 and IPR049063 yielding two nearly identical gene copies of TPS (T2T045g053275/ T2T057g062680) and two highly similar copies (95% identity) of TPP (SCF113g139150/ T2T061g067505). During normal growth, expression of these genes remained low, ranging from 3 to 5 TPM. However, consistent with a role in encystment, TPS and TPP transcripts were strongly induced by galactose treatment (rising over 13-fold from 4.8 TPM to 65 TPM). In contrast, nitroxoline treatment resulted in repression of TPS, while TPP expression was very modestly (3.9 TPM to 8.3 TPM, z-score ∼0.3) induced. Hypoxia yielded modest induction for both TPS and TPP, and only transiently.

Cyst-specific protein 21, another recognized encystment marker in *Acanthamoeba* [55,71], shares homology with universal stress proteins (UspA-like) that are members of the adenine nucleotide alpha hydrolase-like superfamily and are critical for prolonged cellular survival under environmental stress [78]. To identify putative cyst specific proteins in *B. mandrillaris*, we queried protein annotations for USP-like domains using the PFam domain PF00582 and protein homology to *Acanthamoeba* cyst specific protein (XP_004341987.1). This analysis identified 25 unique genes with predicted USP domains (Fig 5A), including four previously annotated as putative cyst-specific proteins in *Acanthamoeba* (T2T077g082940/T2T092g090915, SCF090g133910, SCF130g147345, and T2T012g020125). Consistent with roles during encystment, all four genes were significantly upregulated in galactose specific expression cluster 3 (Fig 2B). Among these, T2T077g082940/T2T092g090915 were transiently upregulated during hypoxia, whereas none of these genes were upregulated in nitroxoline induced cysts. A few predicted genes with weaker similarity to USPs (for example, T2T061g067810/T2T072g078235) yielded expression patterns that were reversed, with enrichment in nitroxoline, relative to galactose. The global transcriptional pattern of nitroxoline and galactose are highly dissimilar, and this analysis of encystment markers reinforces that notion.

**Fig 5.**
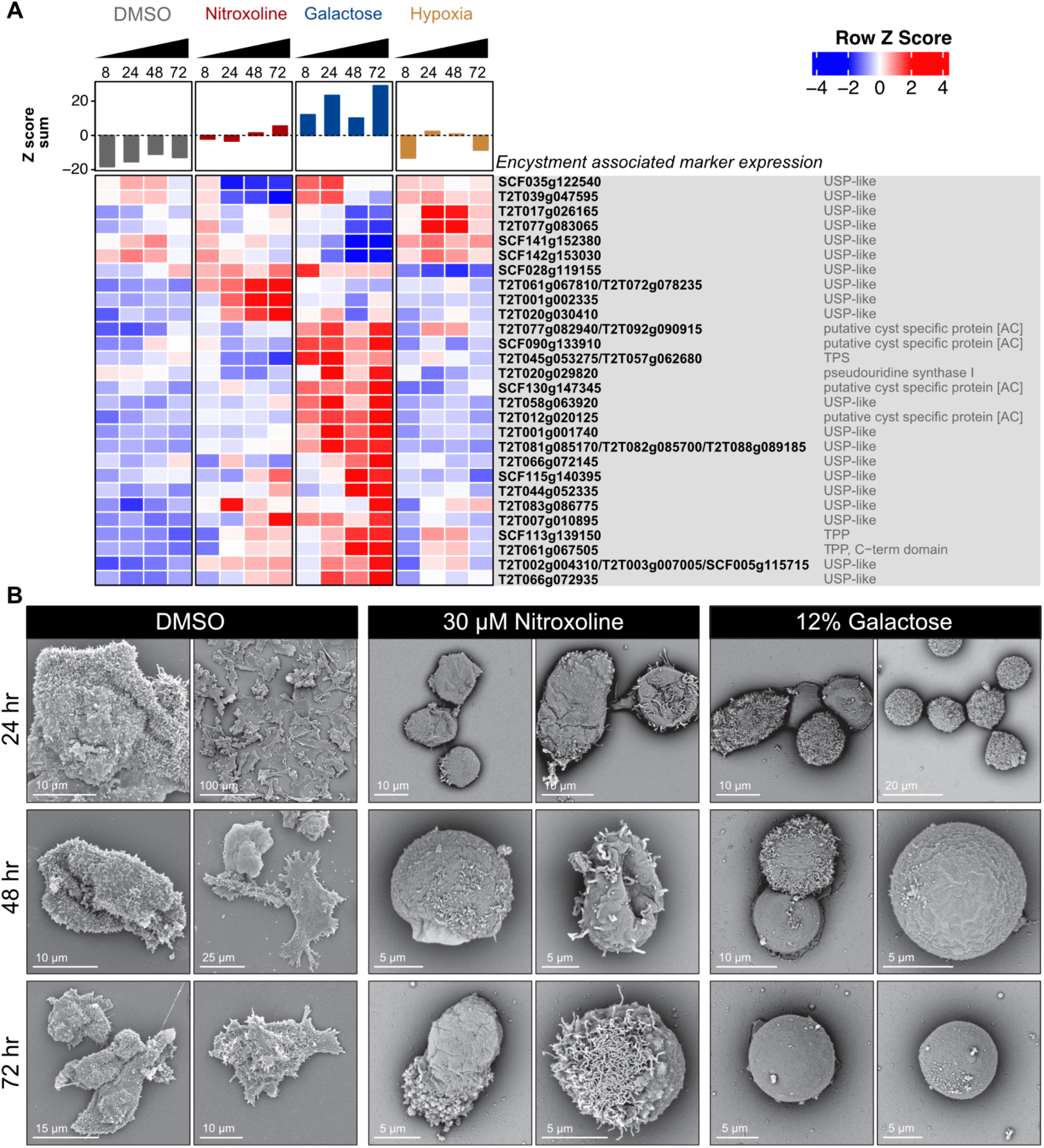
Nitroxoline perturbs the encystment process and undermines structural integrity. (A) Nitroxoline fails to upregulate encystment associated marker genes. Putative trehalose synthesis, universal stress protein family genes (USP-like), and genes sharing homology with *Acanthamoeba castellani* (AC) cyst specific proteins associated with encystment were identified from differentially expressed genes (S1 Table) and their expression patterns were visualized in a heatmap. DESeq2 regularized log transformed counts were averaged for genes sharing at least 0.98 normalized bit score before computing row z scores. (B) Nitroxoline induced cysts show altered morphologies compared to galactose induced cysts. *Balamuthia* trophozoites were treated with 30 μM nitroxoline, 12% galactose, or DMSO control and fixed for scanning electron microscopy at 24, 48, and 72 hours post treatment. Hypoxia was excluded from analysis due to high levels of mixed trophozoite/cyst morphologies.

### Nitroxoline destabilizes the structural integrity of cysts

Scanning electron microscopy (SEM) has been previously used to observe morphological surface features during encystment in free living amoeba [79,80]. Here, SEM (Model Phenom Pharos G2 FEG-SEM, Thermo Fisher, Inc) was employed to qualitatively observe *B. mandrillaris* morphology sampled over a 72-hour time course for DMSO, nitroxoline, and galactose treatments (Fig 5B).

DMSO treated cultures consisted of 100% trophozoites featuring highly pleomorphic morphologies, appearing either rounded, elongated, flattened, or with branching extensions. The surface of *B. mandrillaris* trophozoites was covered with small hair-like projections as previously noted [81] of unknown composition or function. Galactose treated cultures rapidly progress to mature fully rounded, nearly featureless spheres, completely losing their rough polytrichous appearance. Nitroxoline treatment also appears to induce loss of trophozoite features. However, unlike galactose, nitroxoline treated *B. mandrillaris* do not progress to mature cysts but instead appear to arrest with incomplete conversion, often displaying apparent structural defects, including likely extrusions of cytoplasm (Fig 5B). While qualitative, these SEM data suggest nitroxoline initiates a morphological transformation like encystment, but without successful progression. This is also consistent with our previous data showing the morphological changes mediated by nitroxoline are irreversible, unlike galactose induced cysts, as well as the transcriptional comparison between galactose and nitroxoline.

### Nitroxoline prevents recrudescence from galactose induced cysts

Nitroxoline can eliminate *Balamuthia* trophozoites [6] and impedes the formation of viable cysts. However, it remained unclear whether nitroxoline could be efficacious against the quiescent pre-formed cysts, which is an important question when considering treatment of human patients. To investigate this, we treated 3-day old galactose induced cysts for 7 days with 12% galactose, 12.5 µM nitroxoline, or 30 µM nitroxoline. Culture morphology was confirmed pre and post drug treatment via light microscopy (S6 Fig). Cysts were manually counted by hemocytometer and equally distributed onto Vero E6 monolayers. The monolayers were observed daily for recrudescence over a 3-week time course and periodically stained with crystal violet to monitor monolayer destruction (Fig 6A). Galactose induced cysts recrudesced rapidly, consuming 80% of the monolayer in three days and the entire monolayer by six days. In comparison, both 12.5 µM and 30 µM nitroxoline protected the monolayer for 21 days with no evidence of trophozoite reemergence (Fig 6B). These data suggest that nitroxoline disrupts both cyst and trophozoite viability.

**Fig 6.**
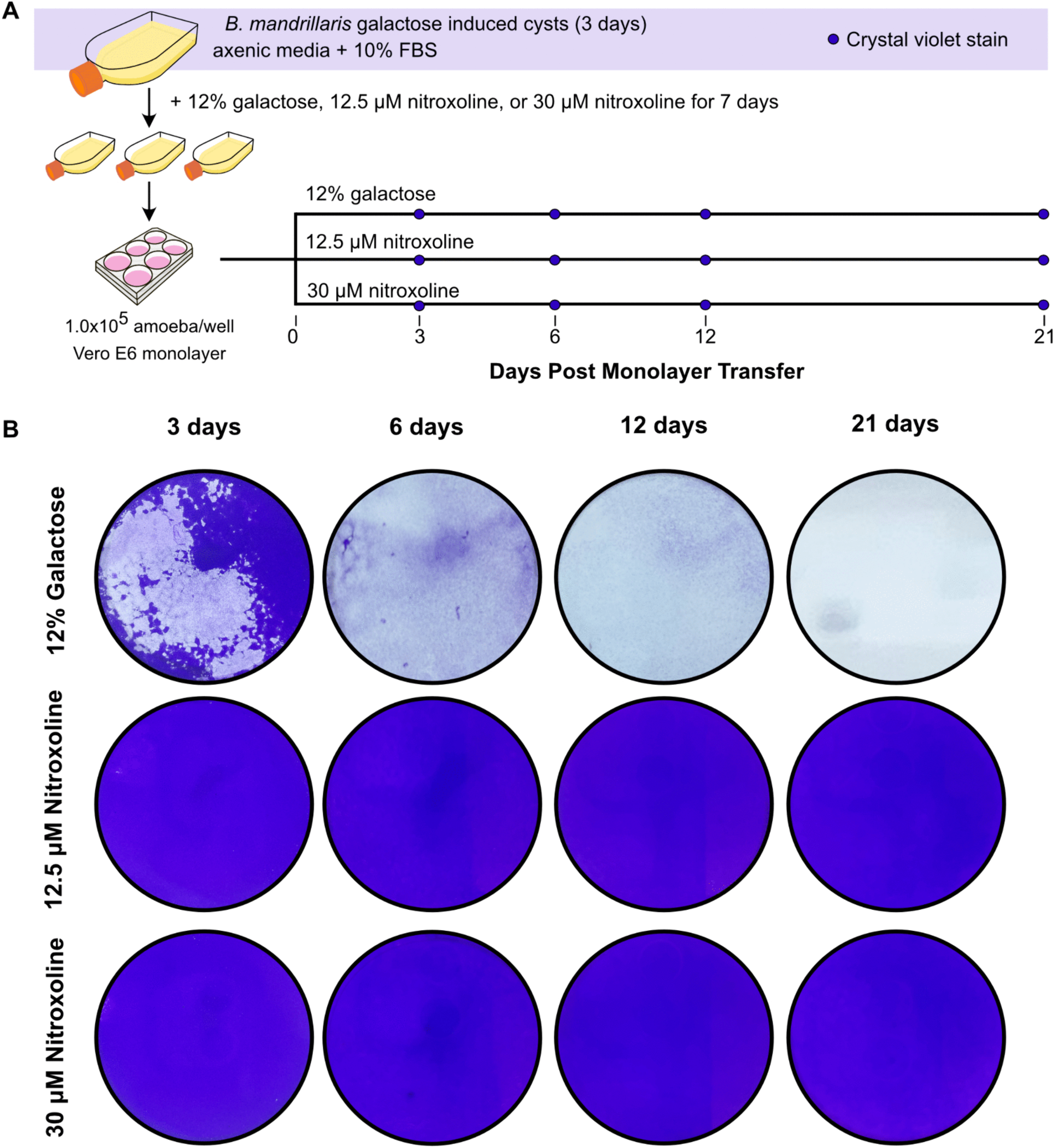
Nitroxoline treatment prevents excystment from galactose induced cysts. (A) Recrudescence assay design. Approximately 25 million *Balamuthia* trophozoites were stimulated with 12% galactose for 72 hours to undergo encystment. Morphology was confirmed via light microscopy (S6 Fig), washed, and transferred to confluent Vero E6 monolayers grown in DMEM. Cultures were observed every day for recrudescence and stained at 3-, 6-, 12-, and 21-days post transfer with crystal violet to visualize monolayer clearance. (B) Representative images of monolayer clearance at indicated times post transfer. Refer to S6 Fig for raw plate images (performed in technical duplicates).

## Discussion

Granulomatous amoebic encephalitis due to *Balamuthia mandrillaris* infection is a devastating, highly fatal condition, with essentially no reliable treatment options, and a mortality rate of 77% even in patients who receive treatment [2]. Due to the small number of cases and high mortality, large scale drug development efforts and traditional randomized control trials of efficacy are not feasible. Our lab previously identified nitroxoline as a promising candidate in a drug repurposing screen. Following subsequent reports of patient recovery after nitroxoline was added to the treatment regimen, nitroxoline has been made available for treatment of *B. mandrillaris* infections in the United States through a CDC expanded access program [16].

To facilitate a mechanistic understanding of nitroxoline’s anti-amoebic activity, we assembled the most complete *Balamuthia mandrillaris* draft genome thus far, produced a plurality of automated annotations for predicted protein sequences and non-coding RNAs, and we provide these as easily accessible .gtf files, downloadable annotations, and a stand-alone genome browser tool. We note that the genome assembly should still be considered a draft, as many loose contigs still exist, with the possibility of mis-assemblies. Regardless, the deliverable of this effort was to provide a foundation for comprehensively identifying coding sequences and non-coding RNAs with high confidence, leveraging the large amount of expression data generated as part of this study. The genome assembly, gene feature definitions, annotations, PacBio HiFi long read data, Illumina short read data, Illumina HiC data, RNAseq data, and random primed metagenomic data are available at NIH BioProject PRJNA1206197. This resource should be invaluable in supporting future functional genomic *Balamuthia mandrillaris* studies, including future genetic manipulations.

This study provides new insights into the mechanism of action for nitroxoline at the transcriptomic, chemical, and structural levels. Treatment with the drug nitroxoline impacts multiple cellular systems, causing a cessation of *B. mandrillaris* growth, catastrophic loss of mRNA, evidence consistent with widespread DNA damage, and impaired or incomplete encystment. Consistent with nitroxoline’s known metal chelating ability, supplementation with copper and iron, but not other divalent metal ions, neutralized its amoebicidal action. Nitroxoline demonstrated efficacy against both trophozoites and cysts, an important consideration given the observation of mixed trophozoite and cyst morphologies in infected tissues. Notably, previous drug screens have not addressed *Balamuthia* pre-formed cyst viability as a parameter.

Impairing functional encystment and preventing excystment represents an attractive therapeutic intervention as potential recrudescence is a cause for concern during clinical treatment of *Balamuthia mandrillaris* infections. Here, nitroxoline was directly compared with two other cellular triggers of encystment, galactose and hypoxia. Galactose induced encystment was associated with transcriptional upregulation of genes encoding putative proteins with known roles in this process in other free-living amoeba, including those with similarity to universal stress domain containing proteins (UspA-like), and the likely trehalose synthesis enzymes TPS and TPP, as well as a nearly complete set of tRNA ligases and associated RNA processing machinery. Hypoxia was a weaker trigger of encystment, with only transient morphological changes, and was associated with transcriptional signatures consistent with low oxygen, including genes encoding putative Hsp20, Hsp70, and Hsp90 family members. Indeed, downregulating Hsp20 in *Acanthamoeba castellani* has been shown to prohibit trophozoites from transitioning into cysts [82]. Comparing transcriptional signatures, both galactose- and hypoxia-induced encystment differed dramatically from nitroxoline treatment.

Given the known ability of nitroxoline to chelate metals, it was not surprising that many cellular functions dependent on or related to metal usage were perturbed, especially those involving copper and iron. Hundreds of proteins are dependent upon copper and iron binding, and we characterized their gene expression in relation to each of the three encystment triggers. Nitroxoline induced encystment showed some overlap in gene expression with hypoxia, mainly with proteins involved in the mitochondrial electron transport chain (especially cytochrome c complex), iron-sulfur cluster binding and assembly, and oxidative stress. These observations are consistent with the fact that many oxygen binding proteins also contain metal binding domains. Furthermore, these findings parallel studies in cancerous cell lines demonstrating that nitroxoline induces oxidative stress, depolarizes mitochondrial membrane potential, and induces mitochondrial-dependent apoptosis [83,84].

Nitroxoline treatment also resulted in a transcriptional response consistent with widespread genomic DNA damage and repair, perhaps due to genotoxic stress associated with the accumulation of replication intermediates or stalled transcriptional complexes, all of which have divalent metal requirements. This genomic crisis was also manifested by catastrophic loss of mRNA content, suggesting a complete loss of pol II transcription competency and/or increased mRNA degradation, an outcome not observed in galactose or hypoxia induced cysts, and a harbinger of irreversible cellular collapse.

Collectively, these data support the notion that nitroxoline sequesters critical iron and copper ions from *B. mandrillaris*, causing widespread metabolic and genomic disruption to normal growth. Given nitroxoline’s well-studied favorable human safety profile, these data further contribute to a rational basis for adding this medication to existing drug regimens for treatment of *B. mandrillaris* infection. Accumulated treatment experience will need to assess real-world clinical efficacy of nitroxoline by comparing survival rates of patients treated with nitroxoline-containing regimens with historical controls. With the addition of nitroxoline to the CDC recommended treatment regimen, and the establishment of a United States national expanded access program, this real-world clinical efficacy question can now be addressed.

## Materials and methods

### Cell culture propagation, encystment, and handling

*Balamuthia mandrillaris* strain CDC:V039 (ATCC 50209) was thawed onto Vero or 293T cell monolayers before passing to modified Cerva’s medium for axenic culture [85]. Routine subculturing of amoeba was performed as previously described by passing 1:3 or 1:6 into fresh axenic medium [6]. Encystment was performed as previously described by incubating amoeba for three days with 12% galactose [6] or by incubating in airtight containers with AnaeroPack™-Microaerophilic gas sachets (Mitsubishi Gas Chemical). When required, amoeba were pelleted between 2500 – 2800 rpm for 5 minutes at room temperature.

### High molecular weight genomic DNA (gDNA) extraction

Approximately 1x10^8^ amoeba were harvested from two HYPERflask vessels (Corning 10031) by gentle tapping and genomic DNA was extracted using Qiagen Genomic Tip 100/G columns. Samples were processed according to manufacturer instructions for cultured cells with the modification of adding 200 ug/ml of RNAase A into Buffer G2 prior to extraction. Samples were incubated in Buffer G2 with RNAse for 15 minutes at room temperature prior to treatment with proteinase K for one hour at 50°C. Sample quality control was assayed via nanodrop, Qubit, and genomic DNA screen tape (Agilent).

### gDNA library preparation for long-read sequencing on a PacBio Sequel IIe

gDNA was aliquoted into two separate halves; one half was sheared with a Covaris g-TUBE, while the other half was size selected using PacBio’s short read eliminator (XL) kit as described by manufacturer. Libraries were prepared according to PacBio’s “Preparing whole genome and metagenome libraries using SMRTbell prep kit 3.0.” Two separate SMRT Cells 8M were sequenced on a PacBio Sequel Ile system (SMRTLink v. 11.0).

### Nextera XT gDNA library preparation

Leftover high molecular weight gDNA was vortexed aggressively for 2 minutes and sheared by pipetting. Sheared DNA was used as input sample for Nextera XT DNA library preparation and prepared according to manufacturer instructions except sample was tagmented for 10 minutes. Library quality was assessed using methods as described above and paired end sequencing of 150 base pair lengths was sequenced on an Illumina NovaSeq X.

### Balamuthia mandrillaris HiC Sequencing

5x10^6^ amoeba (1x T-150) were pelleted and washed in cold PBS. Amoeba were processed according to manufacturer instructions for Phase Genomics Promixo HiC Microbe Kit Protocol v4.5. Sample quality control assessment was done using Qubit dsDNA HS assay and High sensitivity D5000 ScreenTape (Agilent). Paired end sequencing of 75 base pair lengths was conducted on the Illumina NextSeq 2000 P2.

### Draft *Balamuthia mandrillaris* strain CDC:V039 genome assembly

Draft assembly, gene model construction, and automated annotation of coding and non-coding sequences are described in detail in the results section. As described in the results and S1 Fig, our annotation pipeline utilized an arsenal of protein function prediction and annotation tools, consisting of eggNOG, ProteInfer, PANNZER2, and the umbrella annotation suite InterProScan, which encompasses numerous independent analyses. The OpenAI GPT4o large language model was used via the OpenAI API to condense and summarize single line descriptions with NCBI compliant wording using the full collection of annotation descriptions for each putative protein. Single line descriptions with low confidence were flagged for manual review. Product names and descriptions were collapsed by identical genes and close paralogs sharing greater than 0.98 normalized bit scores from a self-aligning DIAMOND run. Proteins without any known functional annotations were relabeled as “hypothetical protein.” The complete set of long reads, assembled contigs, RNAseq reads, HiC Reads, and genome annotations are available at NIH BioProject PRJNA1206197.

### Drug and metal supplemental treatments

All treatments were prepared from powder by dissolving in either 100% dimethyl sulfoxide (DMSO) or tissue culture grade water and filtering with a 0.2 µm syringe. Nitroxoline (Sigma 140325) was dissolved in 100% DMSO to prepare 120 mM stocks, filtered, aliquoted, and stored at -20°C. Deferoxamine was dissolved in water as described by manufacturer (Tocris 5764) at 100 mM, filtered, aliquoted, and stored at - 20°C. Metals (calcium, Sigma C5080; magnesium, Sigma M2773; iron, Sigma F8048; copper, Sigma C8027; manganese, Sigma 416479; and zinc, Sigma Z4750) were prepared fresh prior to each experiment by dissolving in water to prepare 10 mM stocks, filtered, and co-incubated with nitroxoline for five minutes before addition into culture media. All treatments were incubated for three days at 37°C.

### Luciferase assay

Viability was measured using CellTiter-Glo 2.0 reagent (Promega) as previously described [6]. Briefly, 4000 amoebas were seeded into white 96 well plates in 80 ul of axenic media and incubated overnight at 37°C. 5x working stocks of drugs were prepared fresh from freezer stocks and 20 ul of drug treatment was added in 2-fold serial dilutions.

### Scanning electron microscopy

2x10^4^ amoeba were seeded onto collagen coated coverslips (Neuvitro H-12-1.5-Collagen) in a 24 well plate and incubated overnight at 37°C. At specified times, cells were fixed in a solution containing 3% paraformaldehyde (PFA) and 1.5% glutaraldehye (Electron microscopy sciences 16010 and 15714-S) for a minimum of two hours at room temperature. Plates were transferred to 4°C and incubated in fixative overnight before washing and storing in 0.1 mM HEPES for no more than three days prior to sample processing. Samples were dehydrated using serial ethanol washes in 25%, 50%, 75%, and 100% ethanol for five minutes each. Cells were washed a final time with 100% ultra-dry ethanol for two minutes before undergoing critical point drying in a Leica CPD300. Coverslips were sputter coated with 8nm of iridium using a Leica ACE600 sputter coater and imaged on a Phenom pharos G2 Desktop FEG-SEM at 5-10 kV with a mixture of backscatter and secondary-electron detectors.

### RNA extraction for transcriptomics

1.5x10^5^ amoeba were seeded into 6 well plates. Amoeba were treated with either DMSO vehicle control, 12.5 µM or 30 µM nitroxoline, 12% galactose and/or microaerophilic gas generated sachets as described above. RNA was extracted using Direct-zol RNA miniprep kits (Zymo Research) according to manufacturer instructions. RNA integrity was assessed by running the samples on a High Sensitivity RNA ScreenTape (Agilent).

### Poly mRNA enrichment for transcriptomic analysis

Poly(A) mRNA was enriched using the NEBNext poly(A) magnetic isolation module and libraries were prepared for Illumina sequencing with the NEBNext Ultra II Directional RNA library prep kit. Library quality control was assayed using a D5000 ScreenTape (Agilent). Samples were pooled into a single library and paired end sequencing of 147 base pair lengths were sequenced on an Illumina NextSeq 550.

### Total RNA library preparation for integrity assay

ERCCs (ThermoFisher Cat. 4456740) were spiked into total RNA extractions and library preparation was conducted using the NEBNext Ultra II RNA library prep kit for Illumina. Samples were pooled, quality controlled as described above, and paired end sequencing of 150 base pair lengths were sequenced on an Illumina NovaSeq X.

### Recrudescence assay

Five T-150s (∼25 million amoeba) were harvested and stimulated to encyst as previously described with 12% galactose treatment in 20 ml final volume for 72 hours. Morphology was captured via phase microscopy to confirm encystment. Cysts were washed 3x in PBS, split equally into thirds, and resuspended in a T-25 flask with 6 ml of 12% galactose, 12.5 µM nitroxoline, or 30 µM nitroxoline for 7 days. Morphology was captured via phase microscopy to confirm encystment. Cells were washed 3x in PBS, counted via hemocytometer, and 1.0x10^5^ amoeba were seeded onto 6 well plates with technical duplicates on a confluent monolayer of Vero E6 cells. At 3, 6, 12, and 21 days post monolayer transfer, cells were fixed with 3% paraformaldehyde and 1.5% glutaraldehyde for 2 hours and stained for 15 minutes with 1% crystal violet in 20% ethanol. Wells were washed twice with milliQ water before capturing image on light box.

## Data availability

All genomic and RNA sequencing data associated with this project are available at NIH BioProject PRJNA120619. Annotations and transcriptomic expression data may be explored through our stand-alone genome browser tool available at https://github.com/uc-derisilab/derisilab-standalone-balamuthia-browser.git.

## Acknowledgements

For this work, JDR, KAM, AMD, NN, and NS were supported by Biohub, San Francisco. The authors gratefully acknowledge Dyche Mullins of UCSF for SEM access and Samuel Lord of UCSF for SEM training, sample preparation assistance, and imaging. The guidance and expertise in free-living amoeba epidemiology and treatment given by Dr. Julia Haston of the Foodborne, Waterborne, and Environmental Diseases, National Center for Emerging and Infectious Diseases, Centers for Disease Control and Prevention. Heather Stone of the Food and Drug Administration and CURE ID also provided invaluable input on nitroxoline repurposing. Christopher Rice of Purdue University provided advice on experimental design and expertise on amoebic drug discovery studies. Asieris Pharmaceuticals provided nitroxoline pro bono for patient care and conferred on nitroxoline chemistry and use.

## Supporting Information

**S1 Fig.**
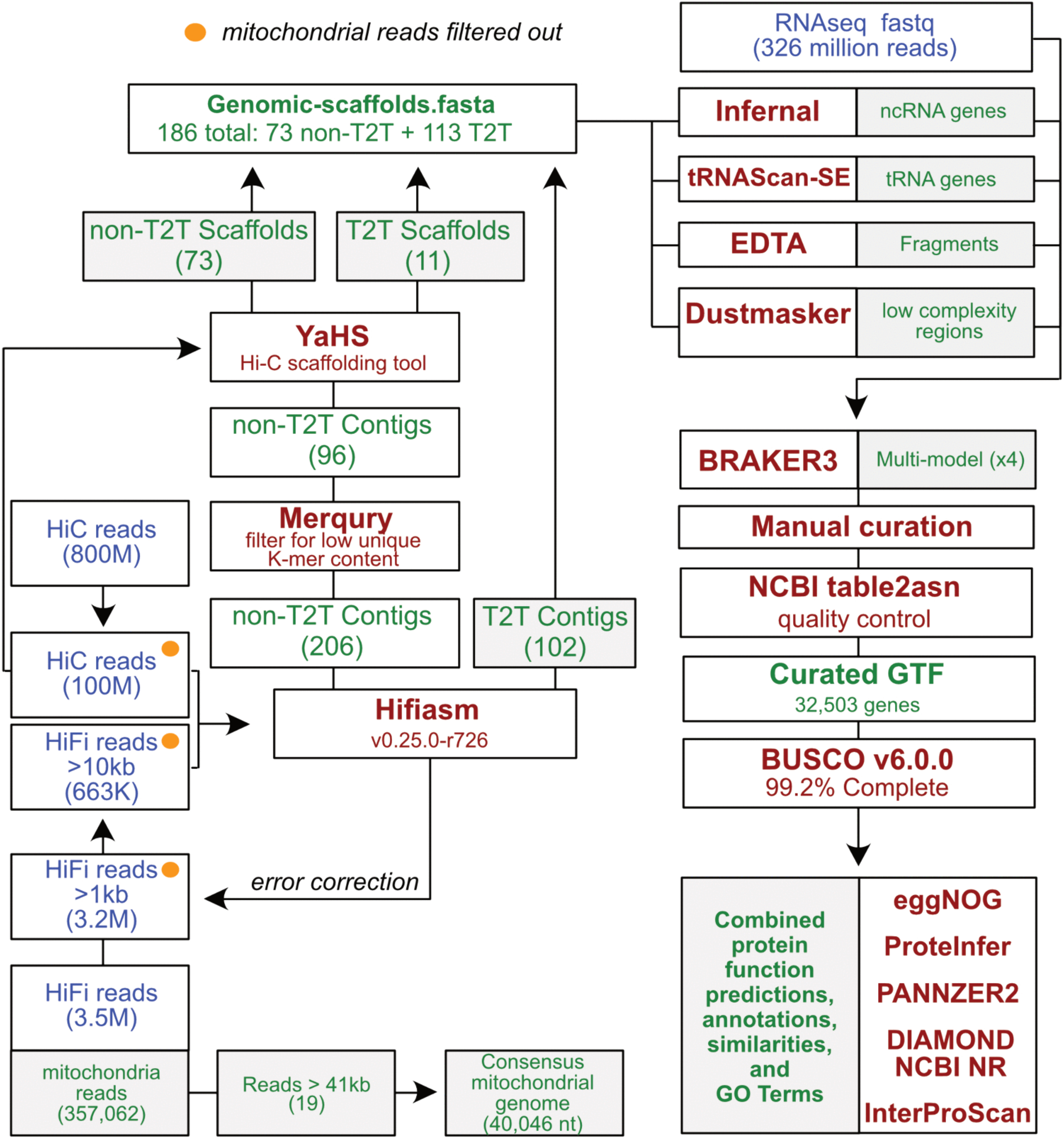
Overview of *Balamuthia mandrillaris* genome assembly and annotation pipeline. PacBio HiFi, Illumina HiC reads, and RNA-sequencing reads were input to the Hifiasm genome assembler. Resulting contigs were evaluated and filtered using Merqury (v1.4.1) to remove contigs with <0.1% unique K-mer contributions, and YaHS (v1.2.2) was used to scaffold non-T2T reads using HiC data. This resulted in a final set of 186 genomic contigs. The contigs were then soft masked for non-coding RNA (ncRNA), transfer RNA (tRNA), mobile elements, and low complexity regions. BRAKER3 (v3.0.8) was then run with the combined RNAseq data using four separate models with softmasking thresholds to provide alternative models for consideration (see S1 Appendix). Gene models were manually inspected and corrected if needed. Annotation of putative coding sequences were processed using a combination of programs, including DIAMOND, eggNOG, ProteInfer, PANNZER-2, and InterProScan, which itself aggregates several annotation modalities.

**S2 Fig.**
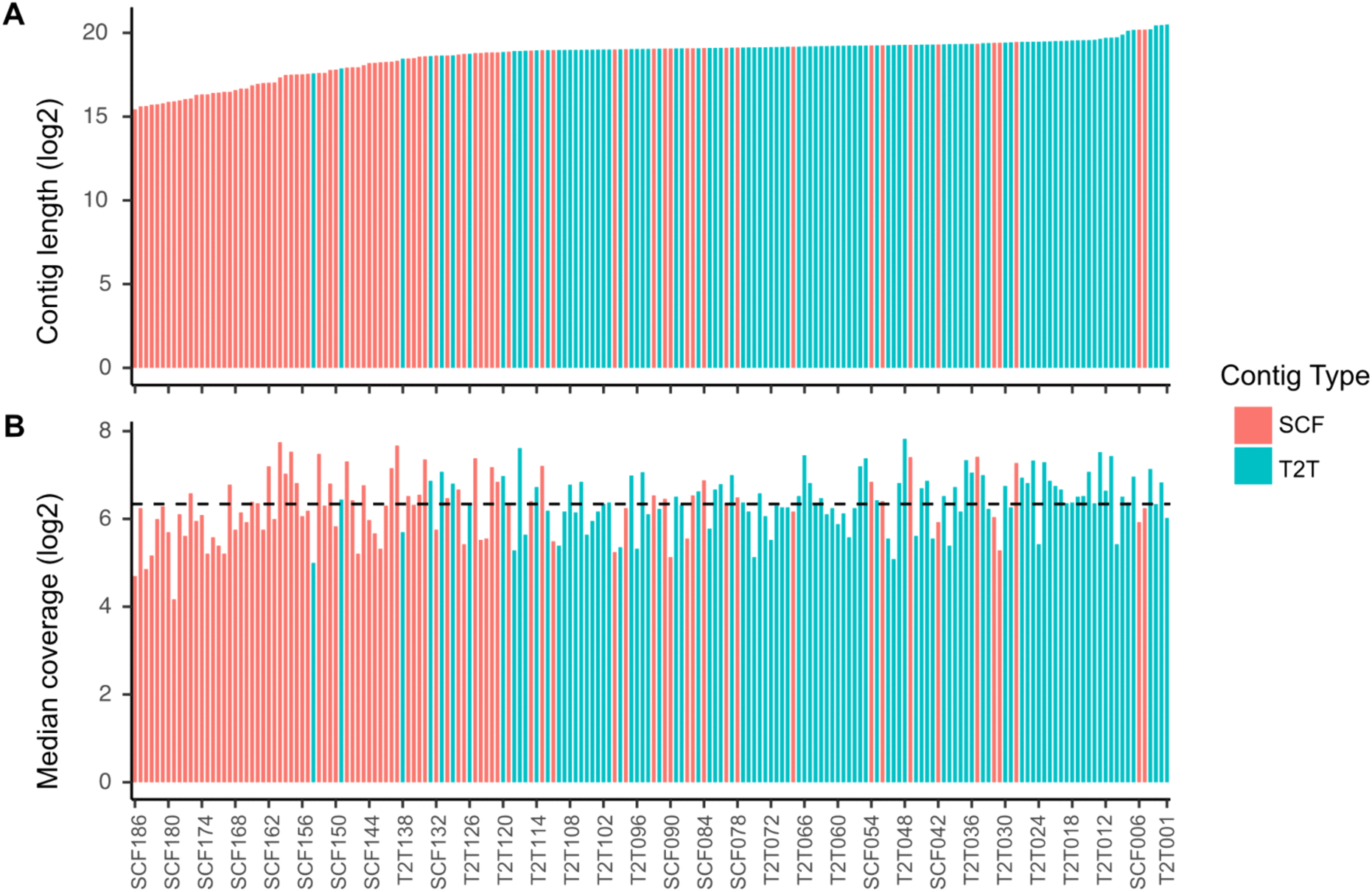
Draft genome assembly contig lengths and median coverage. (A) Log2 transformed length in base pairs for each contig in the *Balamuthia* 186 draft genome assembly, colored by those with telomeric repeats at both ends of the contig (T2T), or scaffold (SCF) contig type, which may have one or no telomeric repeats at the ends. (B) Log2 transformed median coverage depth per contig in draft genome assembly, using only the 10kb or greater read lengths. Black horizontal line depicts average median coverage depth.

**S3 Fig.**
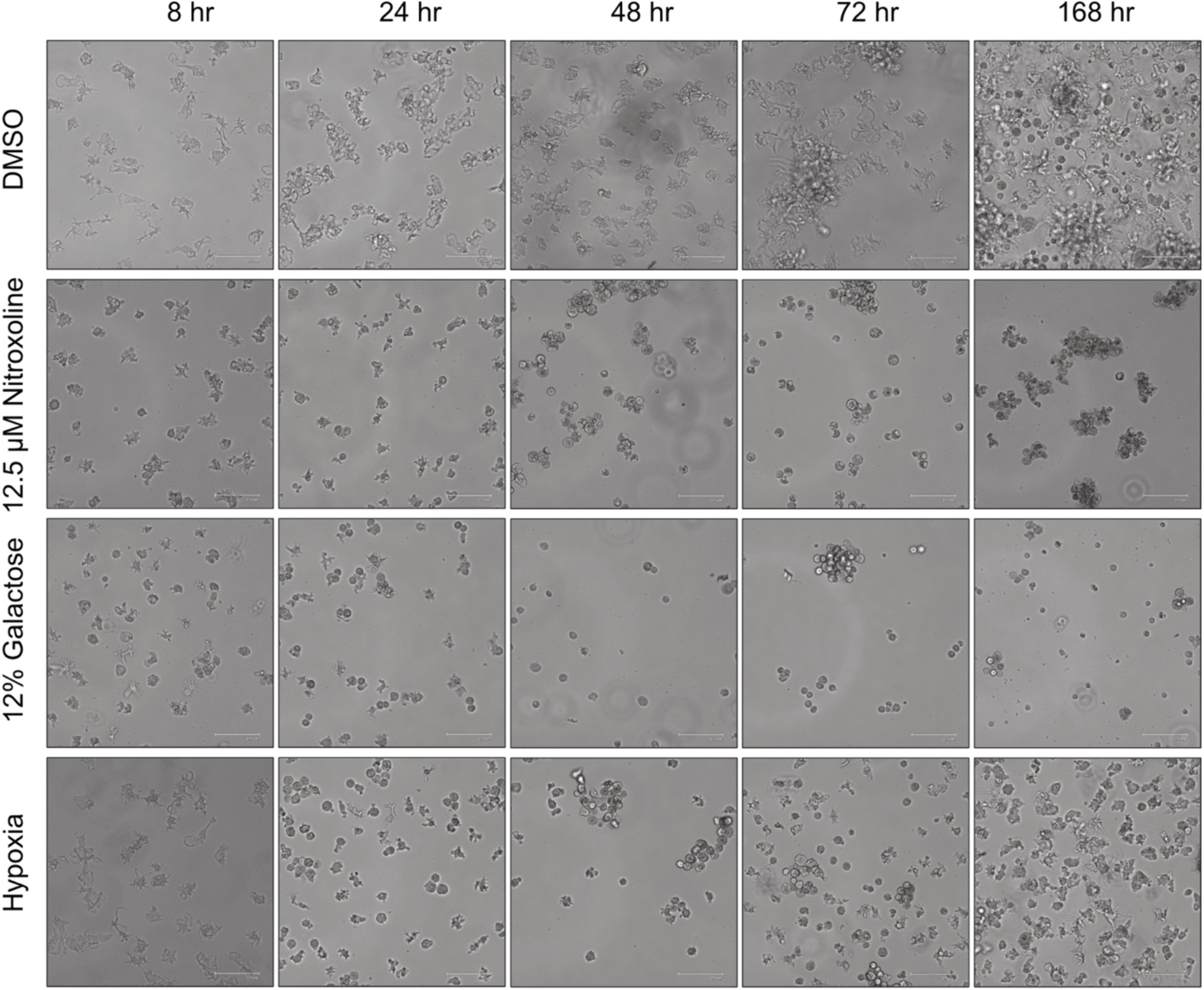
*Balamuthia mandrillaris* morphology in response to encystment. Representative light microscopy images of *Balamuthia mandrillaris* cells taken at time of RNA extraction in response to DMSO vehicle control, 12.5 μM nitroxoline, 12% galactose, or hypoxia treatment.

**S4 Fig.**
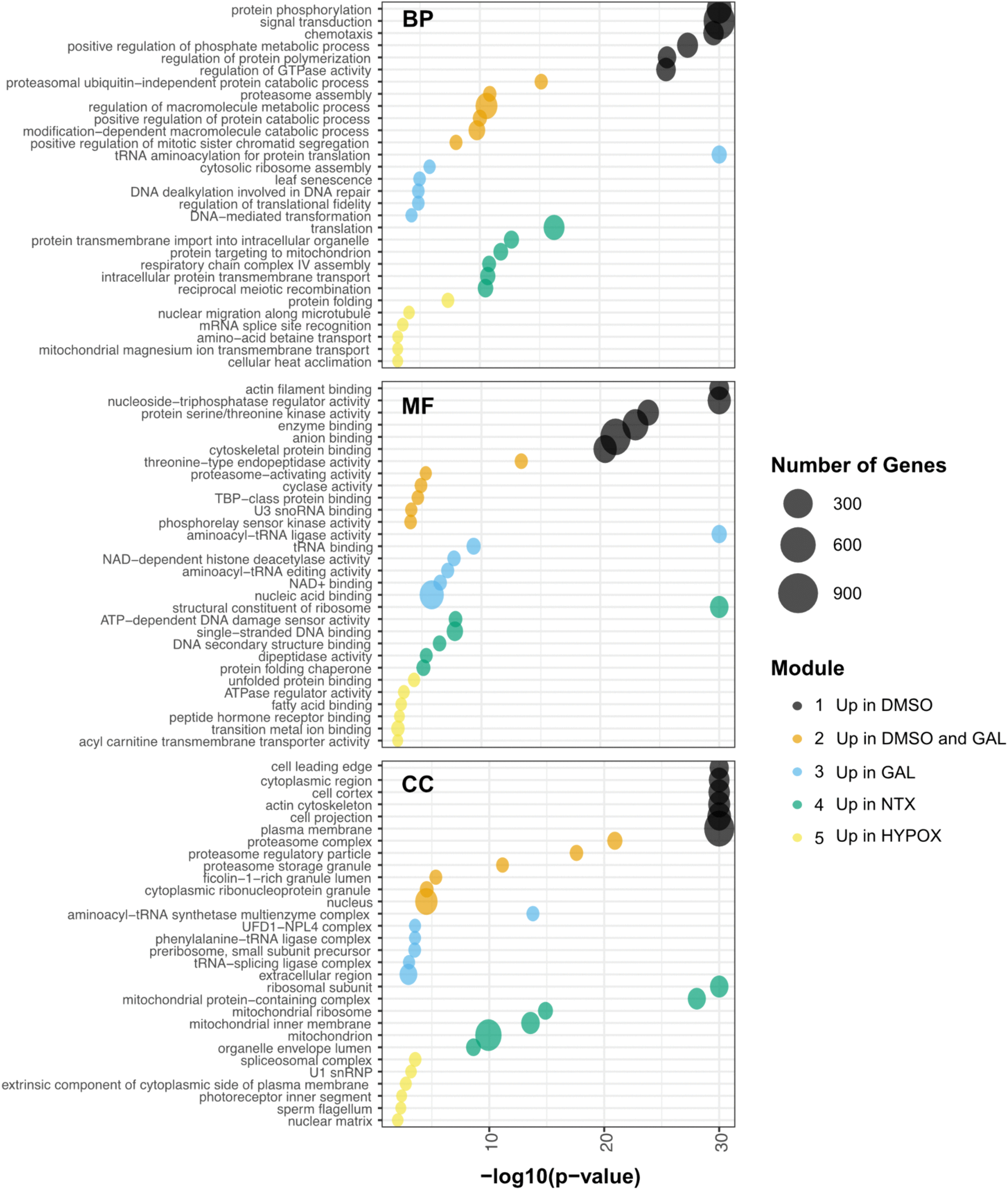
Gene ontology term enrichment of expression clusters of interest. Gene ontology (GO) term enrichment was performed on the five expression clusters of interest highlighted in Fig 2: cluster 1 genes upregulated in DMSO, cluster 2 genes upregulated in DMSO and galactose (GAL), cluster 3 genes strongly upregulated in galactose, cluster 4 genes upregulated in nitroxoline (NTX), and cluster 5 genes strongly upregulated in hypoxia (HYPOX). GO enrichment was performed on biological process (BP), molecular function (MF), and cellular compartment (CC) using the weight algorithm and Fisher exact statistical testing from the topGO R package.

**S5 Fig.**
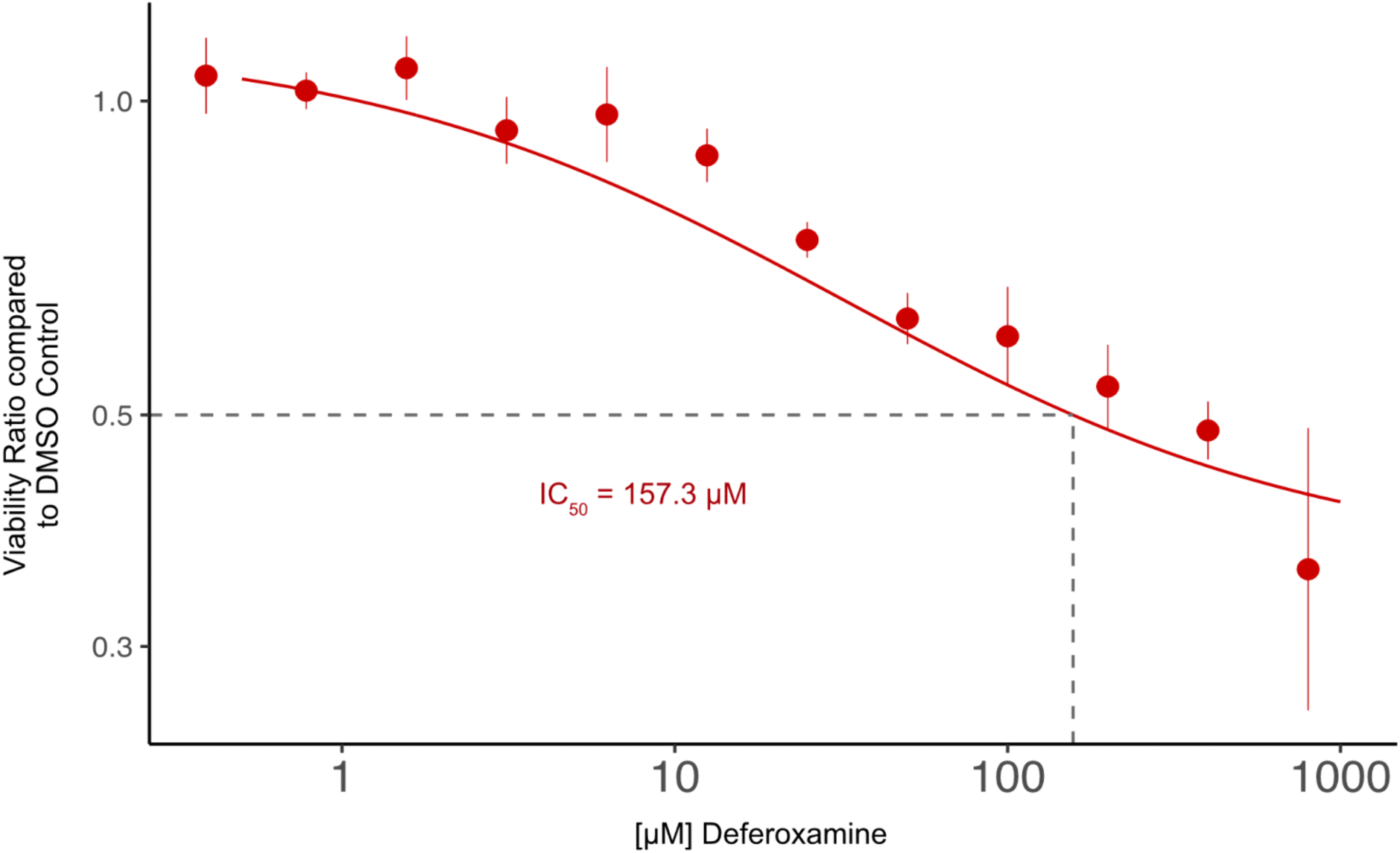
FDA approved iron chelator deferoxamine weakly inhibits *Balamuthia trophozoites*. Deferoxamine or water controls were added in triplicate to trophozoites, and viability was assessed 72 hours later using Promega’s Cell Titer Glo 2.0 reagent. Relative light units (RLU) were normalized to respective water controls (n=3) and the IC50 was determined to be 157.3 μM.

**S6 Fig.**
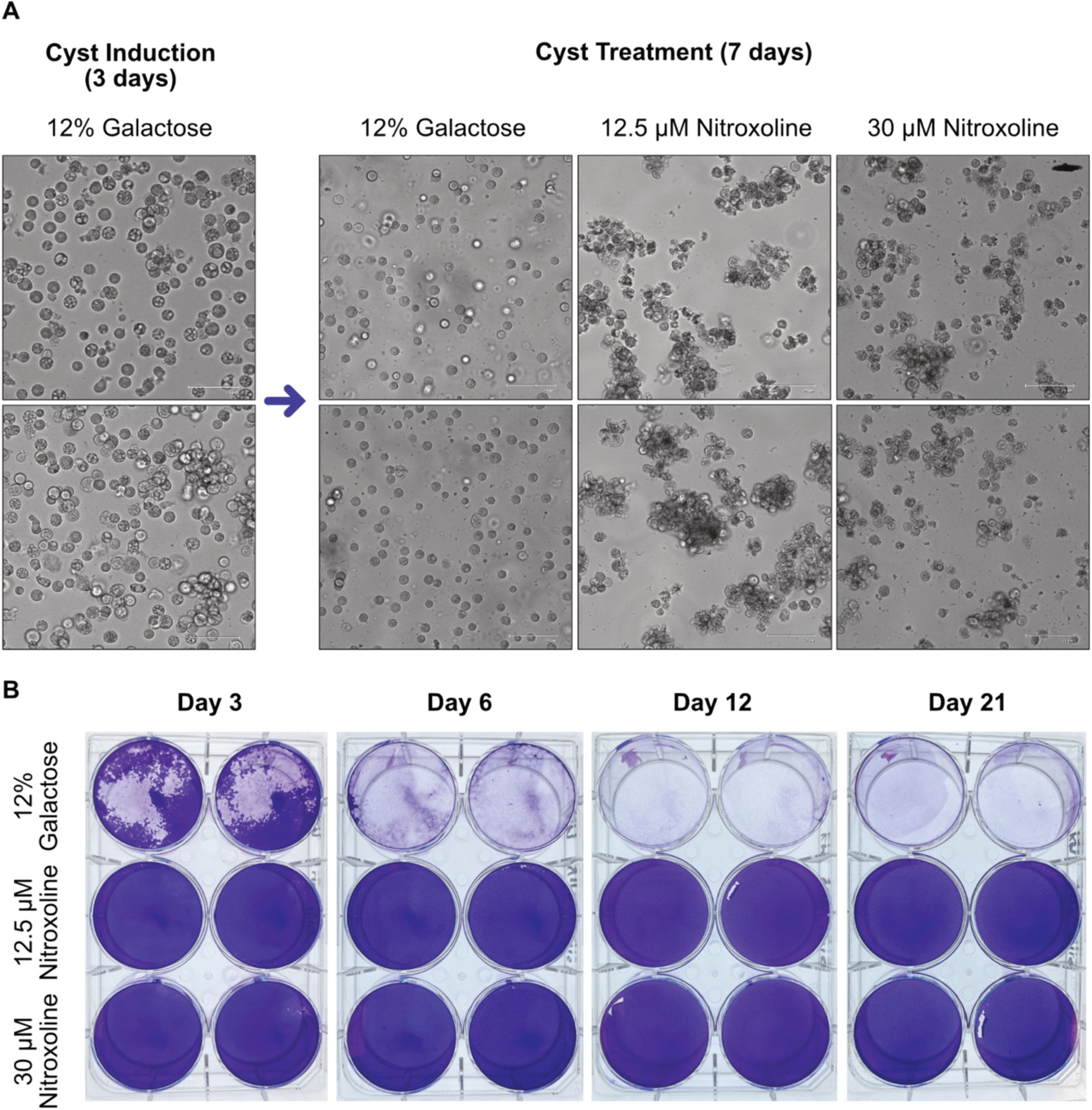
Morphological confirmation of encystment and raw crystal violet stains from recrudescence assay. (A) Morphological confirmation of encystment via light microscopy. Approximately 25 million *Balamuthia* trophozoites were stimulated with 12% galactose for 72 hours to undergo encystment. Morphology was confirmed via light microscopy, and cysts were washed, split equally into three, and incubated with 12% galactose, 12.5 µM nitroxoline, or 30 µM nitroxoline for 7 days. Post treatment, cyst morphology was recorded with light microscopy, washed, and transferred to confluent Vero E6 cell monolayers grown in DMEM. (B) Raw crystal violet stained plates at indicated times post transfer of nitroxoline or galactose induced cysts to Vero E6 monolayers, performed in technical duplicate.

**S1 Table. Row-z score of regularized log transformed counts of 12,328 differentially expressed genes in response to stress induced encystment.**

**S2 Table. Row-z score of averaged regularized log transformed counts of copper, iron, and zinc associated genes upregulated in response to nitroxoline (123 unique genes).**

**S3 Table. Row-z score of averaged regularized log transformed counts of enriched DNA damage and response genes (329 unique genes).**

**S1 Appendix. Excel file containing BRAKER3 model soft masking thresholds, eIF1alpha paralog gene IDs, and curated GO terms for DNA repair and metal associated genes.**

